# Single B cell transcriptomics identifies multiple isotypes of broadly neutralizing antibodies against flaviviruses

**DOI:** 10.1101/2023.04.09.536175

**Authors:** Jay Lubow, Lisa M. Levoir, Duncan K. Ralph, Laura Belmont, Maya Contreras, Catiana H. Cartwright-Acar, Caroline Kikawa, Shruthi Kannan, Edgar Davidson, Benjamin J. Doranz, Veronica Duran, David ER. Sanchez, Ana M. Sanz, Fernando Rosso, Shirit Einav, Frederick A. Matsen, Leslie Goo

## Abstract

Sequential dengue virus (DENV) infections often generate neutralizing antibodies against all four DENV serotypes and sometimes, Zika virus. Characterizing cross-flavivirus broadly neutralizing antibody (bnAb) responses can inform countermeasure strategies that avoid infection enhancement associated with non-neutralizing antibodies. Here, we used single cell transcriptomics to mine the bnAb repertoire following secondary DENV infection. We identified several new bnAbs with comparable or superior breadth and potency to known bnAbs, and with distinct recognition determinants. Unlike all known flavivirus bnAbs, which are IgG1, one newly identified cross-flavivirus bnAb (F25.S02) was derived from IgA1. Both IgG1 and IgA1 versions of F25.S02 and known bnAbs displayed neutralizing activity, but only IgG1 enhanced infection in monocytes expressing IgG and IgA Fc receptors. Moreover, IgG-mediated enhancement of infection was inhibited by IgA1 versions of bnAbs. We demonstrate a role for IgA in flavivirus infection and immunity with implications for vaccine and therapeutic strategies.

## INTRODUCTION

Zika virus (ZIKV) and the four circulating serotypes of dengue virus (DENV1-4) are mosquito-borne flaviviruses with overlapping geographic distributions^1^. Climate change is predicted to further expand the geographic range of mosquito vectors^2–4^, highlighting the need for effective clinical interventions to curb epidemics. The complex antibody response to DENV1-4 has hampered the development of safe and effective vaccines. A first exposure to a given DENV serotype generates potently neutralizing antibodies that typically provide long-term, though sometimes incomplete protection against reinfection by that serotype ^5–7^. However, antibodies that are cross-reactive in binding but not neutralizing activity against other DENV serotypes are also elicited ^8–11^ and pre-existing non-neutralizing antibodies predict the risk of severe disease following secondary exposure to a different DENV serotype ^12–16^. This phenomenon is attributed to a process called antibody-dependent enhancement (ADE), in which non-neutralizing IgG antibodies ^12,17^ facilitate the uptake of bound DENV particles into relevant myeloid target cells via Fc-Fc gamma receptor (FcɣR)-dependent pathways ^18^. ADE-related safety concerns derailed the widespread use of the first licensed DENV vaccine, which increased the risk of severe dengue disease following subsequent infection in previously DENV-naive recipients ^19,20^. As pre-existing IgG antibodies from one prior exposure to ZIKV can also enhance subsequent dengue disease risk^21^, a safe vaccine would ideally induce durable antibodies that can broadly and potently neutralize DENV1-4 and ZIKV.

In contrast to primary DENV exposure, secondary exposure to a different DENV serotype typically elicits broadly neutralizing antibody responses associated with protection against subsequent disease ^8,21–26^. Studying the antibody repertoire in individuals who have experienced multiple DENV infections can thus provide insight into the properties of cross-reactive neutralizing antibody responses that an effective vaccine seeks to mimic. Indeed, a handful of monoclonal broadly neutralizing antibodies (bnAbs) that can potently neutralize DENV1-4 and in some cases, ZIKV, have been isolated from naturally infected individuals living in endemic regions ^27,28,22,29^. The most well-characterized class of flavivirus bnAbs targets a quaternary E-dimer epitope (EDE) spanning both E protein monomers within the dimer subunit ^28,30^. There are two subclasses of EDE bnAbs, of which EDE1 but not EDE2 antibodies can potently neutralize ZIKV in addition to DENV1-4 ^31^. A few antibodies that can cross-neutralize ZIKV and some DENV serotypes have also been described ^32–35^, but other than those of the EDE1 subclass, SIgN-3C is the only known naturally occurring antibody that can potently neutralize ZIKV and all four DENV serotypes ^27,36,37^.

The above antibodies were discovered by sorting hundreds of single B cells from individuals infected with DENV and/or ZIKV, followed by either immortalization or PCR amplification of variable heavy and light chain genes for recombinant IgG production and characterization^38^. Although these approaches have successfully identified bnAbs against many viruses, they are laborious, typically requiring robots and/or large teams to increase throughput. As an alternative high-throughput method, we previously provided proof-of-principle for a single cell RNA sequencing (scRNAseq)-based approach to identify multiple DENV1-4 bnAbs, of which two somatic IgG variants, J8 and J9, were the most potent ^39^. Single cell transcriptomics also allows unbiased profiling of multiple antibody isotypes unlike previous methods, which were largely restricted to isolation of IgG antibodies ^28,33–35,40^.

Here, we have improved upon our scRNAseq-based method to systematically profile the antibody response in 4 individuals whose sera potently cross-neutralized DENV1-4 and ZIKV. We identified 23 new bnAbs, of which a subset displayed neutralization breadth and potency comparable or superior to leading bnAbs in the field but with distinct epitopes. Moreover, one of our newly identified bnAbs neutralized DENV1-4 and ZIKV and is derived from the IgA1 isotype, thus representing the first non-IgG bnAb described against flaviviruses. Notably, monomeric IgA1 versions of newly and previously characterized bnAbs not only retained IgG neutralization breadth and potency, but also inhibited IgG-mediated enhancement of infection in cells expressing both IgG and IgA Fc receptors.

## RESULTS

### Profiling the antibody repertoire of naturally infected individuals

We previously identified bnAbs against DENV1-4 ^39^ from secondary analyses of available scRNAseq data of antibody genes encoded by ∼350 B cells obtained from an unrelated study ^41^. Here, we focused our analysis on B cells from individuals with broadly neutralizing antibody responses to specifically leverage scRNAseq for bnAb discovery (summarized in **Figure 1**). These individuals were enrolled in a prospective cohort in Colombia with confirmed acute DENV or ZIKV infection (**Figure S1**) ^42,43^. We screened longitudinal serum samples from 38 cohort participants for their ability to neutralize prototype DENV1-4 and ZIKV strains in two independent experiments. When tested at a single dilution, no serum sample reproducibly neutralized West Nile virus (WNV), a more distantly related flavivirus included as a control. In contrast, even at the earliest available time point (range: 0 to 7 days after fever onset), serum samples from 26/38 individuals inhibited infection by two or more DENV serotypes by >50% in both experiments (**Figure S1**). This high prevalence of cross-serotype neutralizing activity likely reflects repeated DENV exposures, as confirmed by IgG avidity testing ^42,43^. In addition to broad neutralizing activity against DENV1-4, serum samples from 11/38 individuals reproducibly neutralized >50% infection by ZIKV.

**Figure 1.**
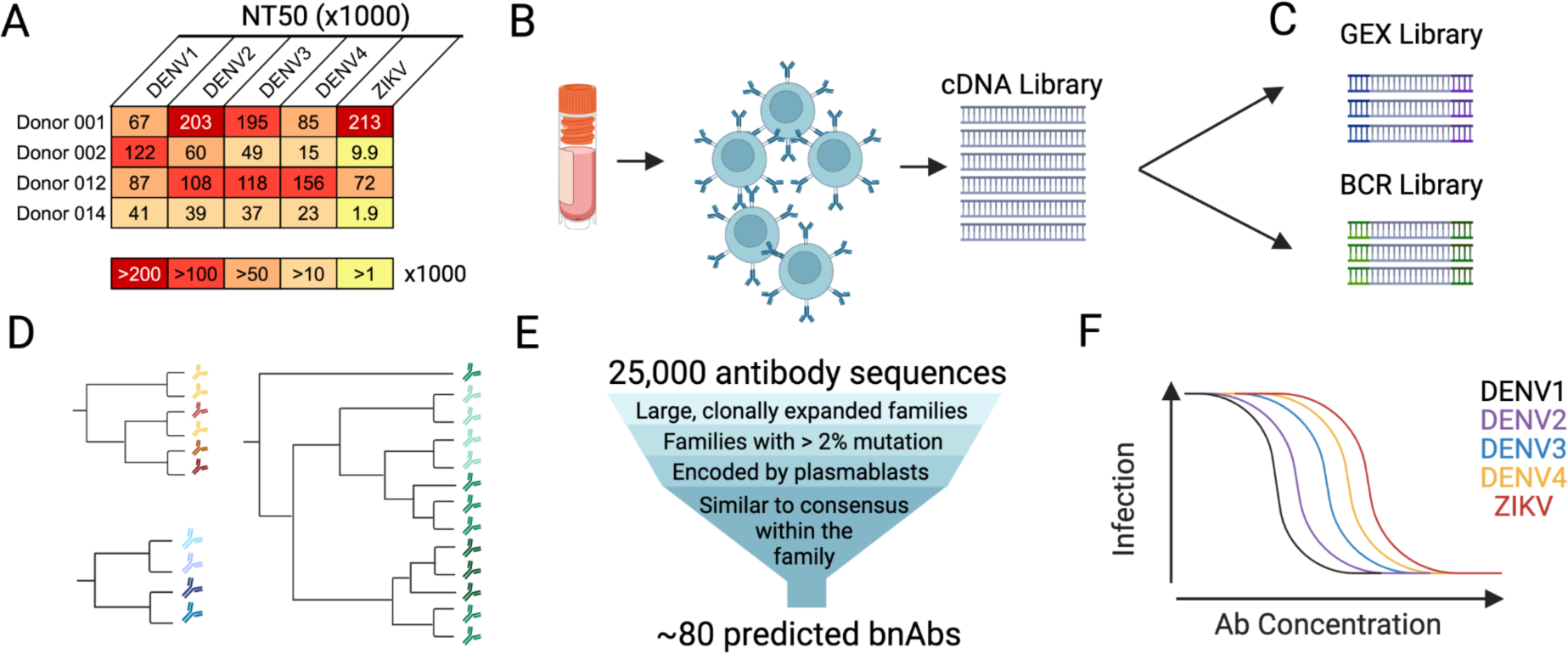
Workflow to identify broadly neutralizing antibodies (bnAbs) from donor samples. **(A)** Serum neutralization profile of 4 cohort participants chosen for downstream analysis based on potent neutralizing activity against DENV1-4 and ZIKV. The mean reciprocal serum dilution that neutralized 50% of virus infection (NT50) in 3 independent experiments is depicted as a heatmap with a darker color indicating greater potency according to the key. **(B)** We isolated B cells isolated from the peripheral blood mononuclear cells (PBMCs) of donors selected in (A) and processed them for **(C)** single-cell RNA sequencing of both global gene expression (GEX) and B cell receptor (BCR)-specific libraries. (**D**) BCR libraries are analyzed by the software package *partis ^51^*, which groups antibodies into clonal families and infers their shared ancestry. **(E)** Antibody sequences most likely to encode flavivirus-specific, high-affinity antibodies are bioinformatically down-selected for functional characterization. (**F)** We recombinantly expressed selected antibodies as IgG1 and screened them for the ability to neutralize DENV1-4 and ZIKV. This figure was created with Biorender.com.

To investigate the properties of broad and potent neutralizing antibody responses, we chose 4 individuals with cross-flavivirus serum neutralizing activity, as confirmed by dose-response neutralization assays (**Figure 1A**). In addition to serum neutralization breadth and potency, these individuals were selected due to the availability of corresponding peripheral blood mononuclear cells (PBMCs) at early time points during which bnAb responses were detected (within 11 days post-fever onset) (**Figure S1**). We chose early time points to maximize our likelihood of detecting transiently circulating plasmablasts. This B cell subset undergoes a large expansion following acute DENV exposure ^25,40,44–47^ and often encodes neutralizing antibodies against multiple DENV serotypes and in some cases, ZIKV, after repeated exposures ^25,27,28,39^. Moreover, unlike memory B cells, plasmablasts constitutively secrete antibodies so their antibody repertoire likely mimics that of contemporaneous serum.

We isolated CD19+ B cells from PBMCs of these 4 donors (**Figure 1B**) for scRNAseq of B cell receptor-specific and overall gene expression libraries (**Figure 1C**). We obtained a total of 25,293 paired antibody coding sequences, with a mean of 6,323 per donor (range 4,644-9,249), comparable to previous studies that profiled antibody repertoires using this method ^48–50^. We first grouped antibodies into clonally related sequences derived from the same rearrangement event (i.e. clonal families, **Figure 1D**) using *partis* ^51^. As shown in **Figure 2A**, the sizes of clonal families and the distributions of B cell subsets within these samples varied substantially. Samples from donors 001 and 012 were dominated by naive B cells that were not members of any clonal family we could discern. By contrast, samples from donors 002 and 014 were composed mostly of plasmablasts in large (4-50 members) or very large (50+ members) clonal families. Antibody isotype distribution also varied by donor: antibodies in samples from donors 001 and 012 were mostly IgM while those from donors 002 and 014 were primarily IgG1 (**Figure 2B**).

**Figure 2.**
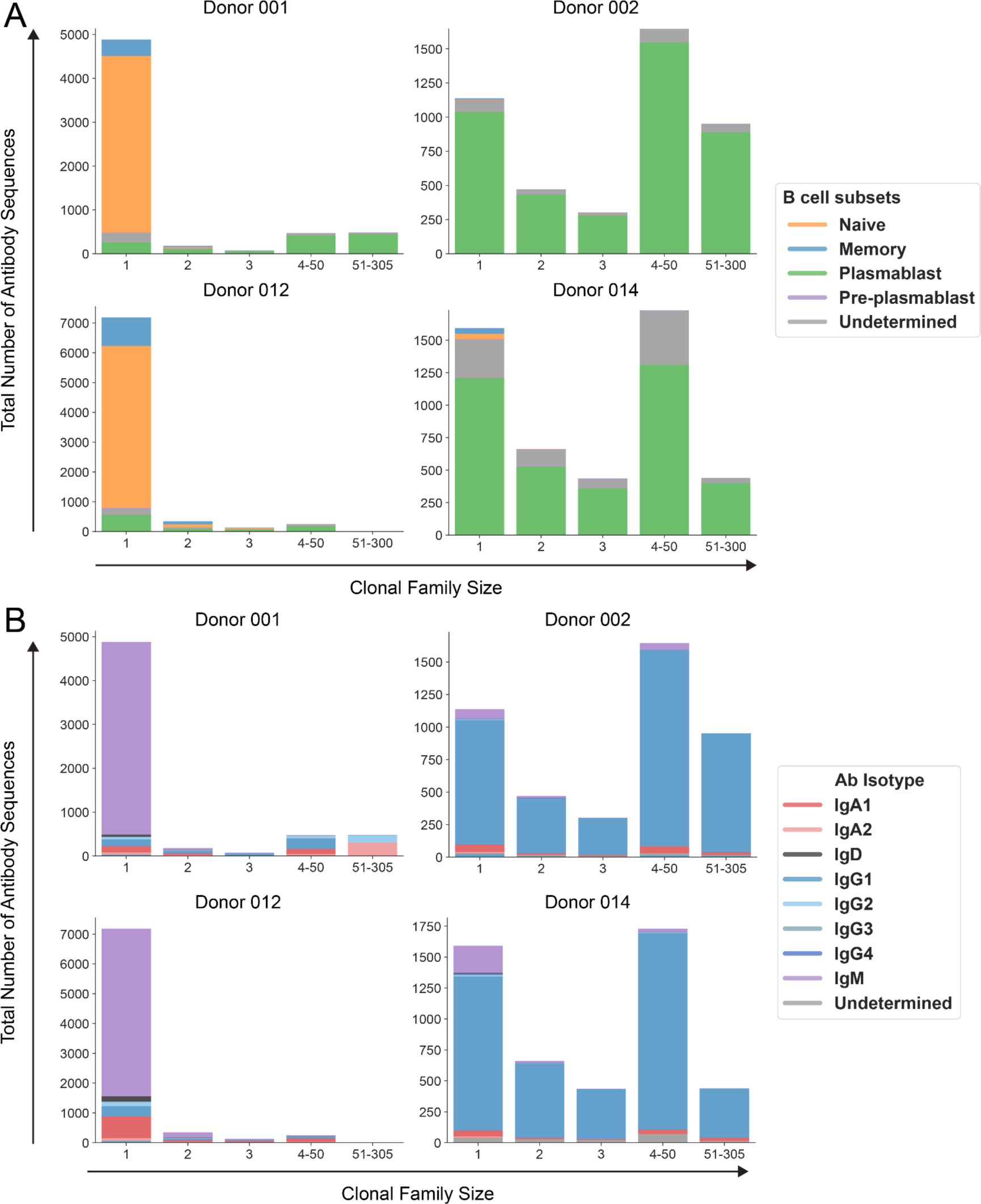
Distribution of B cell subsets and antibody isotypes within clonal families. Graphs depict the number of antibodies encoded **(A)** by distinct B cell subsets and **(B)** as various isotypes in clonal families of different sizes in each of the four donor samples analyzed. B cell subset and antibody isotype were determined by analysis of the cell’s transcriptome as captured by the gene expression library. Only B cells for which a corresponding antibody sequence was observed in the B cell receptor library were included. “Undetermined” B cell subset indicates that that the cell had too few reads or unique molecular identifiers to yield accurate gene expression information as analyzed by 10X Genomics Cell Ranger. “Undetermined” isotype indicates insufficient sequence coverage to determine the constant gene.

### Functional characterization of antibodies

To downselect antibodies for functional characterization, we applied a set of criteria that we and others have found to predict antibody affinity and/or neutralizing activity (summarized in **Figure 1E** and detailed in Methods). Briefly, we chose antibodies that were 1) from clonally expanded families, 2) from families with >2% somatic hypermutation, suggesting antigen-specific selection ^39,50,52^, 3) encoded by plasmablasts as these are often broadly neutralizing ^25,27,28,39^, and 4) most similar to their family’s amino acid consensus sequence, suggesting high affinity ^53^.

We first selected 1-2 antibodies from roughly 20 clonal families per donor to identify families encoding bnAbs. These antibodies were recombinantly expressed as IgG1 by transfection of mammalian cells and the antibody-containing supernatant screened at a single dilution for neutralization of DENV1-4 and ZIKV (**Figure 1F**). As shown in **Figure S2**, the number and neutralization profile of clonal family ‘hits’ varied by donor. For example, of 14 total families tested from donor 001, only two (F05, F07) encoded neutralizing antibodies: F05 antibodies displayed weak ZIKV-specific neutralization, while F07 antibodies neutralized DENV1-3 and ZIKV, but not DENV4. Similarly, only two of the selected families from donor 012 (F12, F15) encoded neutralizing antibodies. In contrast, almost all 26 families from donor 002 neutralized DENV1 and DENV3, though only one (F09) neutralized DENV1-4 and ZIKV. Donor 014 antibodies displayed the broadest neutralization profile: almost all 28 selected clonal families neutralized DENV1-4 and, in some cases, ZIKV with varying potencies. Of these, antibodies from two families (F05 and F09) neutralized DENV1-4 by a mean of 97% and one family (F25) neutralized DENV1-4 and ZIKV by a mean of 92%.

Having identified clonal families encoding bnAbs, we next screened additional members within these families and found that antibodies within a given family generally displayed similar neutralization profiles. For example, all 10 antibodies we selected from family F07 of donor 001 neutralized DENV1, DENV2, DENV3, and ZIKV, but not DENV4. Similarly, all tested antibodies from donor 014 family F09 neutralized all four DENV serotypes but not ZIKV, while those from family F25 broadly neutralized DENV1-4 and ZIKV, though the level of DENV2 and DENV4 inhibition was variable **(Figure S2**). These results demonstrate that our bioinformatics-based approach successfully identified clonal families encoding multiple broadly neutralizing antibodies.

### Identifying antibodies with potent and cross-reactive neutralizing activity

Based on the above crude screens with transfection supernatant, we purified 23 IgG1 antibodies that inhibited DENV1-4 and in some cases ZIKV by >50%. We confirmed their neutralizing activity in dose-response assays and calculated the concentration at which each antibody inhibits 50% of virus infection (IC50). All but one (F15.S01 from donor 012) of these antibodies were from donor 014 and neutralization profiles from dose-response assays were overall concordant with those from the crude screen. We confirmed that the selected antibodies fell into two main categories based on neutralization: 1) those that cross-neutralized DENV1-4 and ZIKV, and 2) those that cross-neutralized DENV1-4 but not ZIKV (**Table S1**).

We compared the neutralization breadth and potencies of our newly identified antibodies to each other and to previously identified bnAbs tested in parallel. EDE1-C10 ^28,31^ and SIgN-3C ^27,37^ represent the only known classes of bnAbs that simultaneously neutralize ZIKV in addition to DENV1-4 (category 1). J9, an antibody we previously isolated from a different donor in the same cohort, potently neutralizes DENV1-4, but not ZIKV (category 2) ^39^. We also included EDE2-A11, which weakly neutralizes ZIKV, unlike EDE1 subclass antibodies ^31^, and MZ4, which neutralizes ZIKV and some DENV serotypes ^33^.

Among all category 1 antibodies tested, the most potent was F25.S02 from donor 014 based on geometric mean IC50 value (**Table S1**). The potency of F25.S02 against ZIKV was comparable to EDE1-C10 (IC50 of 18 and 14 ng/ml, respectively) but was ∼39 times higher than that of SIgN-3C (IC50 of 694 ng/ml). The geometric mean potency of F25.S02 against DENV1-4 was also ∼2-fold higher than that of EDE1-C10 (IC50 of 96 ng/ml versus 207 ng/ml, respectively). Family F25 contained 3 other antibodies that broadly neutralized DENV1-4 and ZIKV. These antibodies (F25.S03, F25.S04, F25.S06) neutralized DENV1, DENV2, DENV3, and ZIKV with relatively similar potency as F25.S02, but they were less potent against DENV4 (IC50 of ∼ 1 μg/ml).

Among the newly identified category 2 antibodies, F09.S05 was most potent; its geometric mean IC50 against DENV1-4 was comparable to the previously identified J9 ^39^ (36 ng/ml and 33 ng/ml, respectively). Additional high-ranking category 2 antibodies include others from family F09 and antibody F05.S03 from family F05.

Thus, we identified several neutralizing antibodies with similar or better breadth and potency compared to existing bnAbs. Even within the same donor, these bnAbs were derived from multiple germline genes and did not display unusually high levels of somatic hypermutation (**Table S2**), as has been reported for some bnAbs against other viruses ^54,55^. For subsequent detailed characterization, we chose the top-ranking antibody from each clonal family of donor 014, namely F25.S02, F09.S05, and F05.S03. **Figure 3A** shows representative dose-response neutralization assays demonstrating that these new bnAbs are roughly as potent, and in some cases, more potent, than previously published bnAbs (**Figure 3B** and **Table S1**).

**Figure 3.**
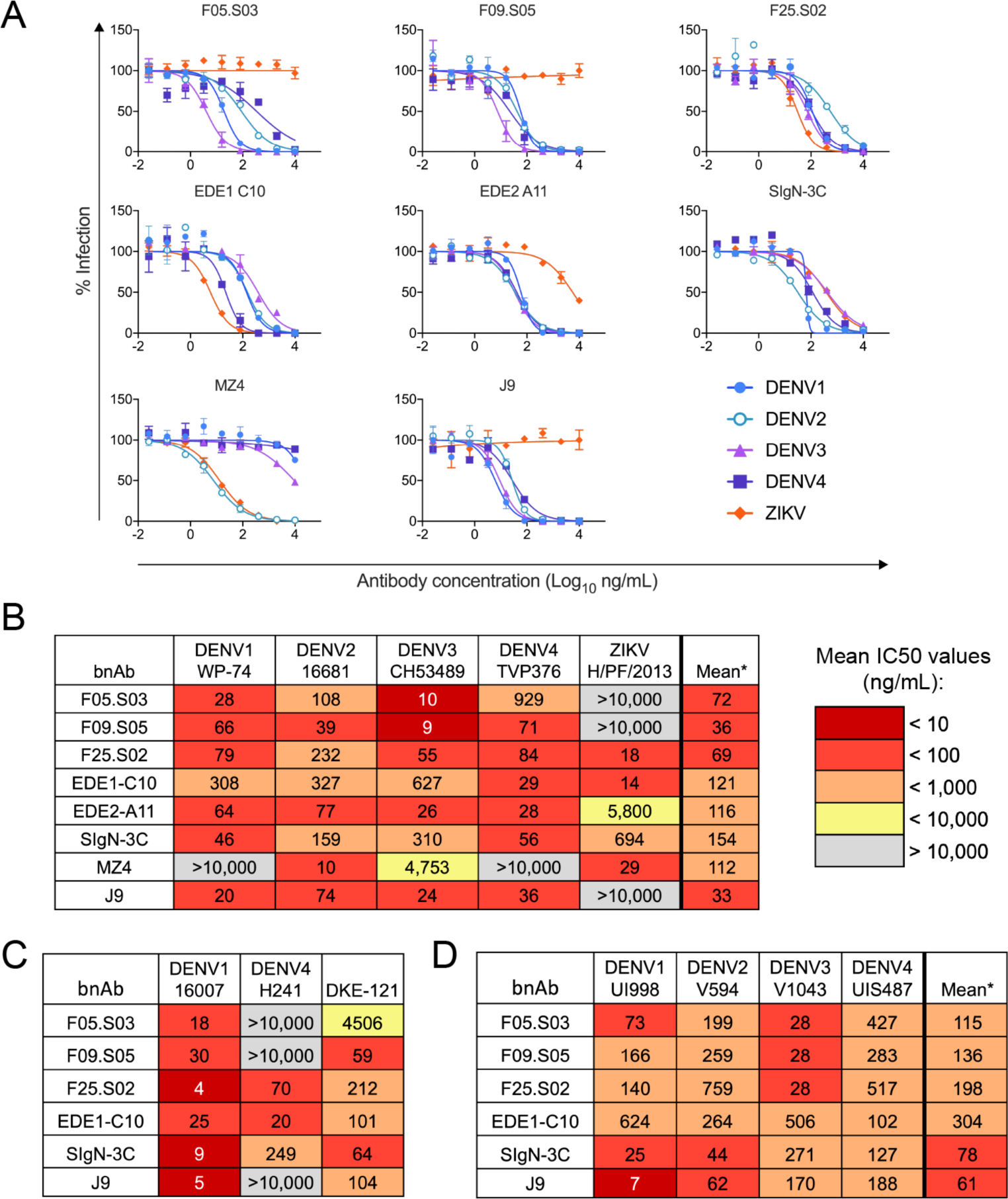
Neutralization profile of top bnAbs expressed as IgG1. **(A)** Representative dose-response neutralization curves of each antibody against the indicated reporter virus particles performed in at least 3 biological replicates in duplicate wells. The data points represent the mean and the error bars represent the range of the duplicates. **(B)** Mean IC50 values for antibody-virus pairs shown in (A) and compiled from Table S1. *The final column displays the geometric mean IC50 values against neutralized viruses. **(C)** IC50 values against additional DENV variants selected due to known antigenic divergence from the panel in (A). Values shown are means from at least two biological replicates. **(D)** Mean IC50 values against fully infectious DENV clinical isolates from 2004-2006. The values are means of at least two biological replicates. *The final column displays the geometric mean IC50 of each antibody against the four viruses. In (B-D), IC50 values are displayed as heatmaps according to the key. Gray indicates that 50% neutralization was not observed at the highest antibody concentration tested (10,000 ng/ml).

### Newly identified antibodies neutralize flavivirus antigenic variants

There is antigenic variation even within a given DENV serotype ^56–58^, which is composed of distinct genotypes ^59,60^. For example, the DENV1 strain West Pac-74 (WP-74) used in the above screens belongs to genotype IV, which is the most antigenically distinct within this serotype ^57^. Additionally, this DENV1 strain is thought to display altered structural dynamics that globally affect antigenicity ^61,62^. To rule out the possibility that DENV1 inhibition was limited to an unusually neutralization-sensitive strain, we confirmed that our novel bnAbs also potently neutralized the genotype II DENV1 strain 16007 (IC50 range of 4 to 30 ng/ml, **Figure 3C**). DENV4 also displays antigenic variation across genotypes (I and II) that circulate in humans ^63,64^. Many of our newly identified lower-ranking antibodies and some known bnAbs neutralized the DENV4 genotype II TVP376 strain used in the above screens with modest potency (**Figure S2**). When tested against the DENV4 genotype I strain H241, we found that category 1, but not category 2 bnAbs retained neutralization potency **(Figure 3C)**. This preferential neutralization of DENV4 genotype II by most antibodies is consistent with previous observations ^64–67^. We also tested our bnAbs against DKE-121, a recently identified strain that is so distantly related to existing serotypes that some have proposed a fifth serotype ^68–70^. Excitingly, F25.S02 and F09.S05 potently neutralized DKE-121(IC50 of 212 and 59 ng/ml, respectively), though F05.S03’s neutralization of this strain was relatively weak (IC50 of 4500 ng/ml) (**Figure 3C**).

Except for DKE-121, most strains used above were lab-adapted and isolated many decades ago (1956 - 1982). Additionally, most were tested as single-round infectious reporter virus particles (with the exception of H241, which was tested as a replication competent virus). Reassuringly, F25.S02, F09.S05, and F05.S03 also neutralized more contemporaneous, fully infectious DENV1-4 clinical isolates collected between 2004 and 2007 with geometric mean IC50 values lower than for the known bnAb EDE1-C10 but higher than SIgN-3C and J9 (**Figure 3D**).

Aside from genetic diversity, flavivirus antigenic variation can also arise from heterogeneous virion maturation states resulting from inefficient cleavage of prM, a chaperone for the E protein. Many but not all flavivirus-specific antibodies preferentially neutralize incompletely mature virions that retain prM on the surface ^71–73^. Importantly, there is increasing evidence that the ability to neutralize the structurally mature form of flaviviruses is important for *in vivo* protection ^74,75^. We tested the ability of our novel bnAbs to neutralize either partially mature DENV2 or ZIKV produced under standard conditions or more fully mature viruses produced in the presence of excess furin to enhance prM cleavage ^72^ (**Figure S3**). As controls, we included antibodies E60 and ZV-67, which poorly neutralize mature forms of DENV2 and ZIKV, respectively, resulting in a large fraction of non-neutralized virions even at high antibody concentrations, consistent with previous studies ^72,76,77^. In contrast to these control antibodies, whose IC50s against mature virus is 15-30 fold higher than against partially mature virus (**Figure S3**), F25.S02, F09.S05, and F05.S03 potently neutralized DENV2 regardless of maturation state (maximum IC50 fold change of 2.7). Moreover, F25.S02 was *more* potent against the mature form of ZIKV (15-fold decrease in IC50). We also observed preferential neutralization of mature ZIKV by known bnAbs EDE1-C10 and SIgN-3C. Overall, these results demonstrate that our new bnAbs can neutralize flavivirus antigenic variants arising from both genetic and structural heterogeneity that are relevant for vaccine efficacy ^66,67,75^, though the ability to broadly neutralize multiple DENV4 genotypes was restricted to F25.S02.

### Mapping E protein determinants of antibody binding

Many potently neutralizing flavivirus antibodies target complex epitopes displayed optimally on virions and not on soluble monomeric E protein ^78^. To determine the E protein oligomeric form recognized by our bnAbs, we performed ELISA to assess binding to soluble monomeric E protein or to virus particles of the prototype DENV2 16681 strain. Unlike antibody B10, which we previously showed to efficiently bind E proteins displayed in both contexts ^39^, F25.S02, F09.S05, and F05.S03 bound efficiently to E proteins displayed on virus particles only, similar to the known bnAb EDE1-C10 ^28^ (**Figures 4A-B**). These results suggest that our newly identified bnAbs preferentially recognize quaternary epitopes.

**Figure 4.**
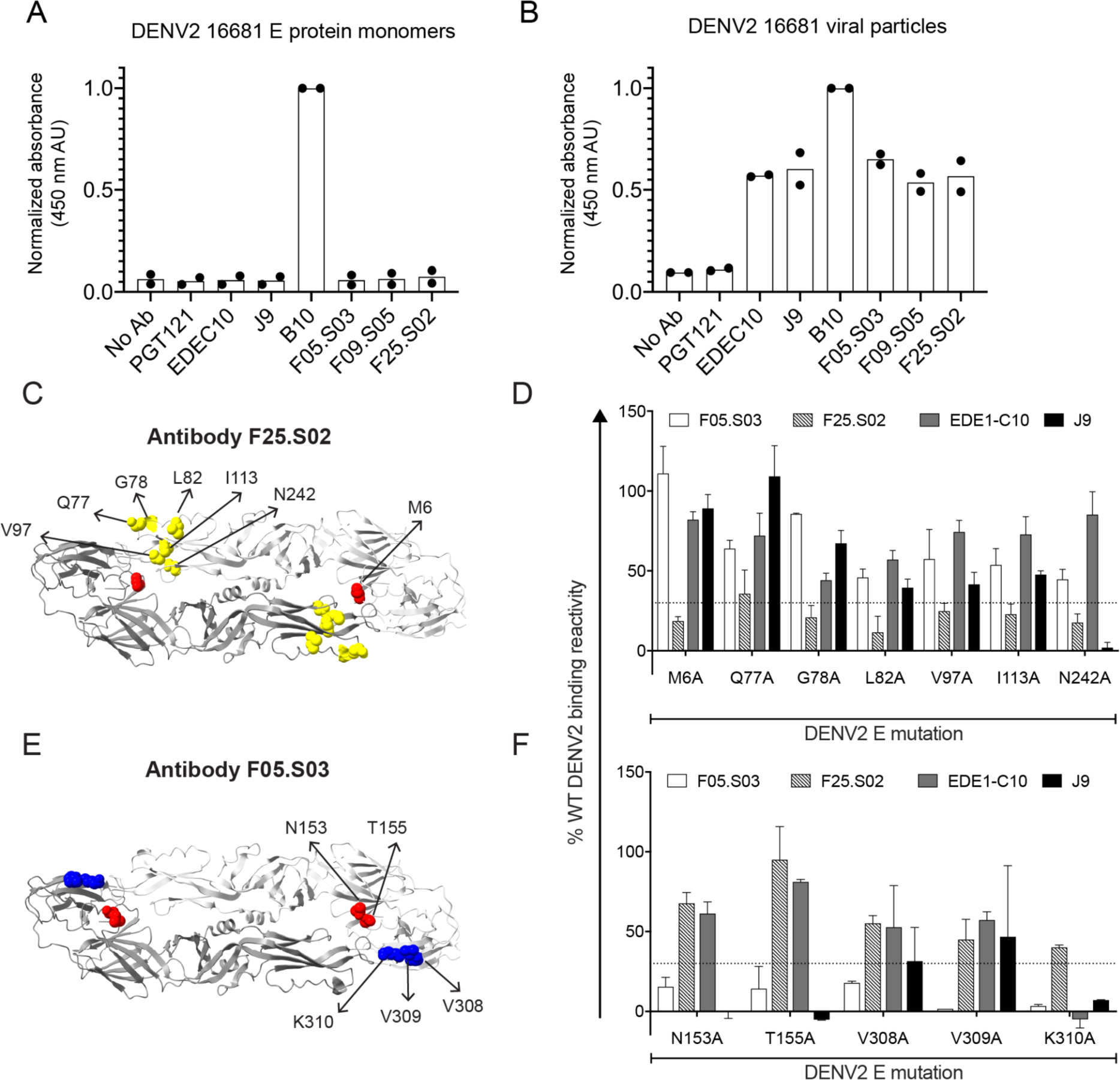
Determinants of E protein binding by bnAbs. Relative binding efficiency (measured by ELISA) by the antibodies indicated to **(A)** E protein monomers (**B**) or virus particles of DENV2 16681. The results are from two independent experiments, each performed in duplicate wells. The absorbance of each duplicate, reported in arbitrary units (AU), was normalized to the wells that received positive control antibody B10 ^39^. HIV-specific antibody, PGT121 ^133^ was used as a negative control. Data points represent the normalized means of each experiment and the bars represent the means of the two experiments. **(C-F)** DENV2 16681 E protein sites important for binding by antibody **(C)** F25.S02 or **(E)** F05.S03 are shown on the ribbon structure of the DENV2 E dimer (PDB: 1OAN) and labeled on one monomer. Sites in E domains I, II, and III are depicted in red, yellow, and blue, respectively. Bar graphs show the mean binding reactivity to individual alanine mutants that selectively impact **(D)** F25.S02 or **(F)** F05.S03 as a percentage of wildtype (WT) DENV2 E protein reactivity. Binding of control antibodies EDE1-C10 and J9 to these mutants was tested in parallel. Error bars show the range of binding reactivity from two independent experiments. The dotted line indicates 70% reduction in antibody binding activity to mutant compared to WT.

To identify E protein amino acid residues critical for binding, we screened antibodies against a shotgun alanine-scanning mutagenesis library of DENV2 prM/E proteins ^39,79^. As controls, we included known bnAbs EDE1-C10 and J9. We identified alanine mutations that specifically reduced F25.S02 or F05.S03 binding by >70% relative to wild type DENV2 (**Figures 4C-F**; **Table S3** shows screen results against the entire library).

For F25.S02, all E residues identified as important for binding were located in domain II (G78, L82, V97, I113, N242) with the exception of M6 in domain I (**Figures 4C-D**). Mutation at these residues minimally impacted binding by the known bnAb EDE1-C10, which retained 50-85% of wild type binding reactivity (**Figure 4D**). EDE1-C10 and F25.S02 are further distinguished by their dependence on K310A, which abolished binding by EDE1-C10, but not by F25.S02 (**Figure 4D**). Thus, although F25.S02 and EDE1-C10 display a similar neutralization profile against DENV1-4 and ZIKV, their binding determinants on DENV2 are distinct.

For F05.S03, mutation at E residue N153 or T155 in domain I, each of which abolishes a potential N-linked glycosylation site, reduced binding efficiency by ∼85% (**Figures 4E-F**). The presence of this potential N-linked glycosylation site has also been shown to be important for recognition by J9 ^39^(**Figure 4F**) and by the EDE2 subclass of bnAbs ^28^. A shared feature of these antibodies is potent neutralization of DENV1-4, but not ZIKV (**Figure 3**). Other residues important for F05.S03 binding include V308, V309, and K310 in E domain III. Of these, K310A also strongly reduced binding efficiency by J9 (**Figure 4F**).

Despite testing multiple conditions (data not shown) we did not detect binding of F09.S05 to wild type DENV2 in this format. Thus, we used an alternative mapping approach (below) for this antibody.

### Mapping neutralization determinants

As F09.S05 neutralized DENV1-4 but not ZIKV, we screened neutralizing activity against a previously described DENV2 library encoding mutations at solvent accessible E residues that were identical or similar across representative DENV1-4 strains but different from ZIKV ^39^. Specifically, amino acids at these E protein sites in DENV2 16681 were substituted with corresponding ZIKV H/PF/2013 amino acids individually or in combination to identify those that reduce antibody potency against DENV2 and thus comprise the neutralization epitope. We also tested a subset of DENV2 alanine mutations identified in the binding screen above to validate their role in neutralization.

Except for the K310A mutation in E domain III, which reduced F09.S05 potency by ∼14-fold, mutations that strongly impacted F09.S05 neutralizing activity were in domain I (**Figures 5A**). Removing the potential N-linked glycosylation site through mutation at residue N153 or T155 abrogated neutralization, while the nearby V151T mutation reduced F09.S05 potency by ∼50-fold. Combining V151T with H149S abolished neutralizing activity. The glycosylation site mutations also abolished neutralization by F05.S03 (**Figure 5B)** and J9 (**Figure 5C**), consistent with results from our binding screen above (**Figure 4F**) and our previous study with J9 ^39^. In addition to these shared residues important for neutralization, we identified unique determinants that distinguished F09.S05 and F05.S03 from each other and from the previously characterized J9. For example, although the individual S145A and H149S mutations minimally impacted F09.S05 and J9 (maximum of 5-fold change in IC50), each mutation reduced F05.S03 neutralization potency by ∼20-fold. Moreover, the combination of K47T+F279S mutations in domain I minimally impacted F09.S05 and F05.S03 (<4-fold IC50 change, **Figures 5A-B**), but reduced J9 potency by 76-fold (**Figure 5C**).

**Figure 5.**
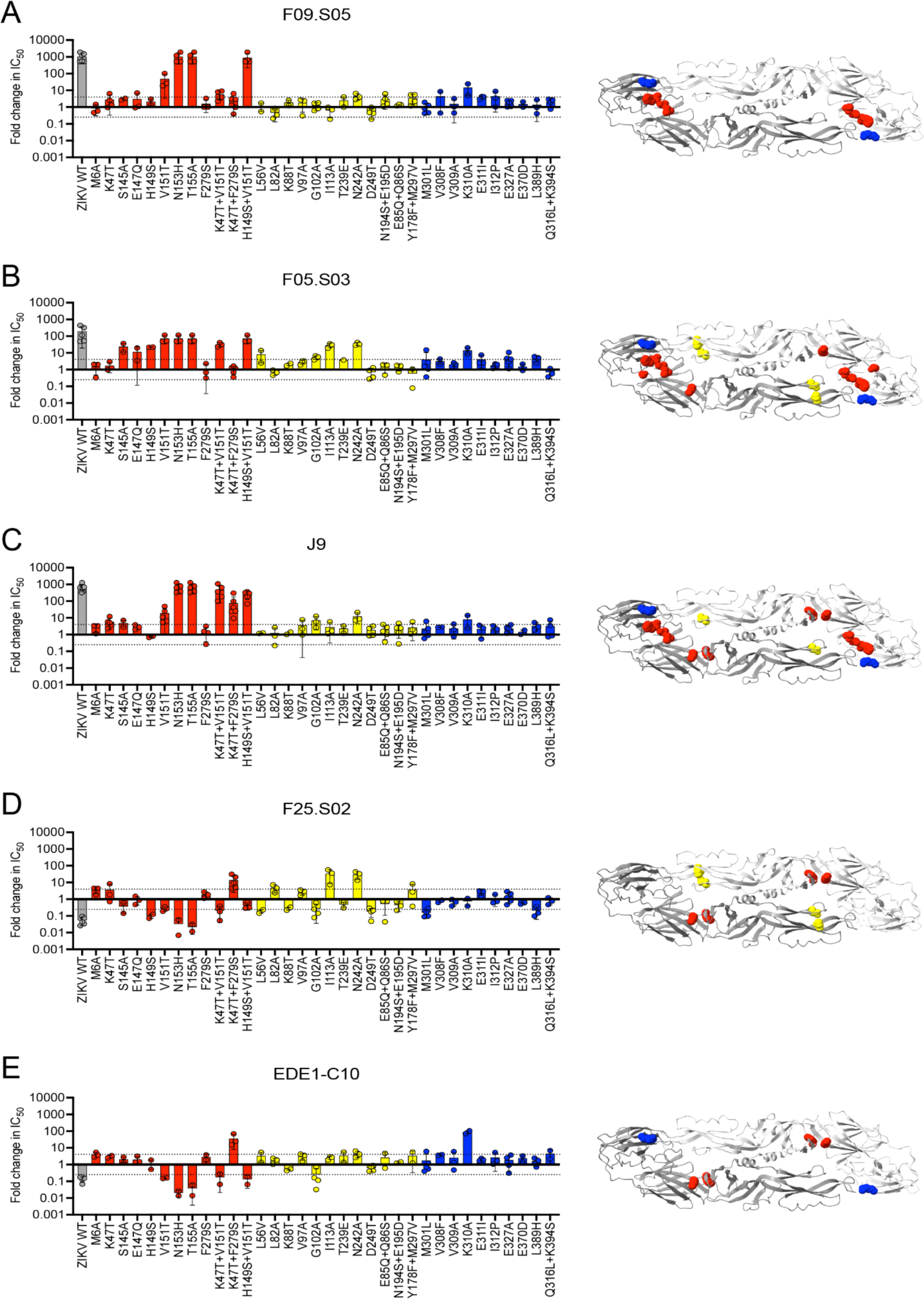
E protein residues critical for neutralization by bnAbs. (Left panel) Bar graphs show the mean IC50 fold change against DENV2 16681 reporter virus particles encoding E protein variants relative to wild type (WT) DENV2 for antibodies **(A)** F09.S05, **(B)** F25.S02, **(C)** F05.S03, **(D)** EDE1-C10, and **(E)** J9. Values of 1, >1, and <1 indicate no change, decreased sensitivity, and increased sensitivity of mutant relative to WT DENV2, respectively. Mean values were obtained from at least 2 independent experiments shown as individual data points in which WT and mutant DENV2 were tested in parallel. WT ZIKV H/PF/2013 (gray) was included as a control. Error bars indicate the standard deviation (n>2) or range (n=2). In each graph, the dotted horizontal line represents a 4-fold IC50 change. (Right panel) For each bnAb, sites of mutations that reduced neutralization potency when tested either individually or in combination by > 4-fold are depicted as spheres on both monomers of the DENV2 E dimer subunit (PDB 1OAN). Sites in E domains I, II, and III are shown in red, yellow, and blue, respectively.

As mentioned, the N153 and T155 glycosylation site mutants abolished neutralization by F09.S05, F05.S03, and J9, which neutralize DENV1-4 but not ZIKV. In contrast, when tested against EDE1-C10 and F25.S02, both of which neutralize ZIKV in addition to DENV1-4, these mutations *increased* neutralization potency by up to 50-fold (**Figures 5D-E**). Another shared feature between EDE1-C10 and F25.S02 is a reduced neutralization potency against the K47T+F279S double mutation in E domain (36- and 14-fold IC50 increase, respectively). However, there were distinct neutralization determinants for these bnAbs. Specifically, the I113A and N242A mutations in domain II each reduced F25.S02 potency by ∼30-fold (**Figure 5D**) but minimally impacted EDE1-C10 neutralization (<4-fold IC50 change, **Figure 5E**). Conversely, the K310A mutation in domain III strongly reduced EDE1-C10 (∼90-fold IC50 increase, **Figure 5E**) but not F25.S02 potency (0.7-fold IC50 change, **Figure 5D**). These results are consistent with the alanine binding screen (**Figure 4D**). Thus, despite some similarities, we identified E residues that distinctly impact neutralization by newly discovered bnAbs relative to each other and to known bnAbs.

### Effect of antibody valency on neutralizing activity

To gain insight into the epitope arrangement on virions, we compared the neutralization potency of F25.S02, F09.S05, and F05.S03 tested as bivalent IgG or monovalent Fab against DENV2 and ZIKV (**Figure S4**). Except for F09.S05, the Fab versions of all antibodies tested, including known bnAb controls, EDE1-C10 and SIgN-3C, failed to neutralize DENV2 by at least 50% at the highest antibody concentration tested (400 nM), suggesting that bivalent engagement is important for potent DENV2 neutralization by these antibodies ^80^. Although SIgN-3C IgG neutralized ZIKV with moderate potency, no neutralization was detected with Fab, consistent with previous findings ^37^. In contrast, EDE1-C10 and F25.S02 retained the ability to completely neutralize ZIKV as Fab. Although IgG versions of EDE1-C10 and F25.S02 neutralized ZIKV with similar potency, their Fab neutralization profiles were more distinct; unlike EDE1-C10 Fab, which retained relatively potent neutralization consistent with previous findings (<10-fold increase in IC50 compared to IgG) ^80^, F25.S02 neutralized ZIKV with much reduced potency as Fab (64-fold increase in IC50 compared IgG). These results suggest that EDE1-C10, SIgN-3C, and F25.S02 target distinct epitopes on ZIKV.

### Neutralizing activity of IgA1 antibodies is similar to or better than IgG1 versions

As neutralizing activity is traditionally thought to be dependent mainly on changes within the antibody variable region, neutralizing antibodies have typically been tested as the IgG1 subclass, regardless of their native isotype ^81^. Moreover, most studies profiling the neutralizing antibody repertoire against flaviviruses have specifically isolated IgG antibodies ^28,33–35,40,44^. While we did not bias our scRNAseq-based approach towards a particular antibody isotype, we initially expressed and screened all antibodies as IgG1, similar to previous studies. Given increasing evidence that antibody Fc isotype can impact neutralizing activity against many viruses ^82–86^, we used scRNAseq data to confirm that the native isotype of almost all 23 antibodies downselected for detailed characterization was indeed IgG1 (**Table S2**). However, unlike other flavivirus bnAbs described here or previously, our top-ranking bnAb, F25.S02 was derived from the IgA1 isotype.

To investigate the impact of isotype on neutralizing activity, we expressed F25.S02, EDE1-C10, and SIgN-3C as monomeric or dimeric IgA1 and compared their neutralization profile to IgG1 versions. Although we purified IgA1 dimers by size-exclusion chromatography (SEC), we could not exclude the presence of higher order polymers ^87^ by SDS-PAGE analysis **(Figure S5**) so we refer to these antibodies as polymeric IgA1 hereafter. Monomers that were produced in transfections lacking a J chain expression plasmid appeared identical to monomers that were separated from polymers via SEC (**Figure S5**), but for simplicity all experiments were performed using the former.

As shown in **Figure 6**, all 3 bnAbs retained neutralization breadth and potency as monomeric IgA1. Moreover, while F25.S02 monomeric IgA1 and IgG1 displayed comparable potency against DENV1-4 and ZIKV (maximum of 2-fold IC50 change), monomeric IgA1 versions of EDE1-C10 and SIgN-3C were more potent against some viruses (**Figure 6B**). For example, compared to their IgG1 versions, EDE1-C10 and SIgN-3C monomeric IgA1 antibodies were ∼4 times more potent against DENV3, though sample sizes (n=3) were too small to achieve statistical significance. SIgN-3C potency against ZIKV was also 9 times higher as monomeric IgA1 compared to IgG1.

**Figure 6.**
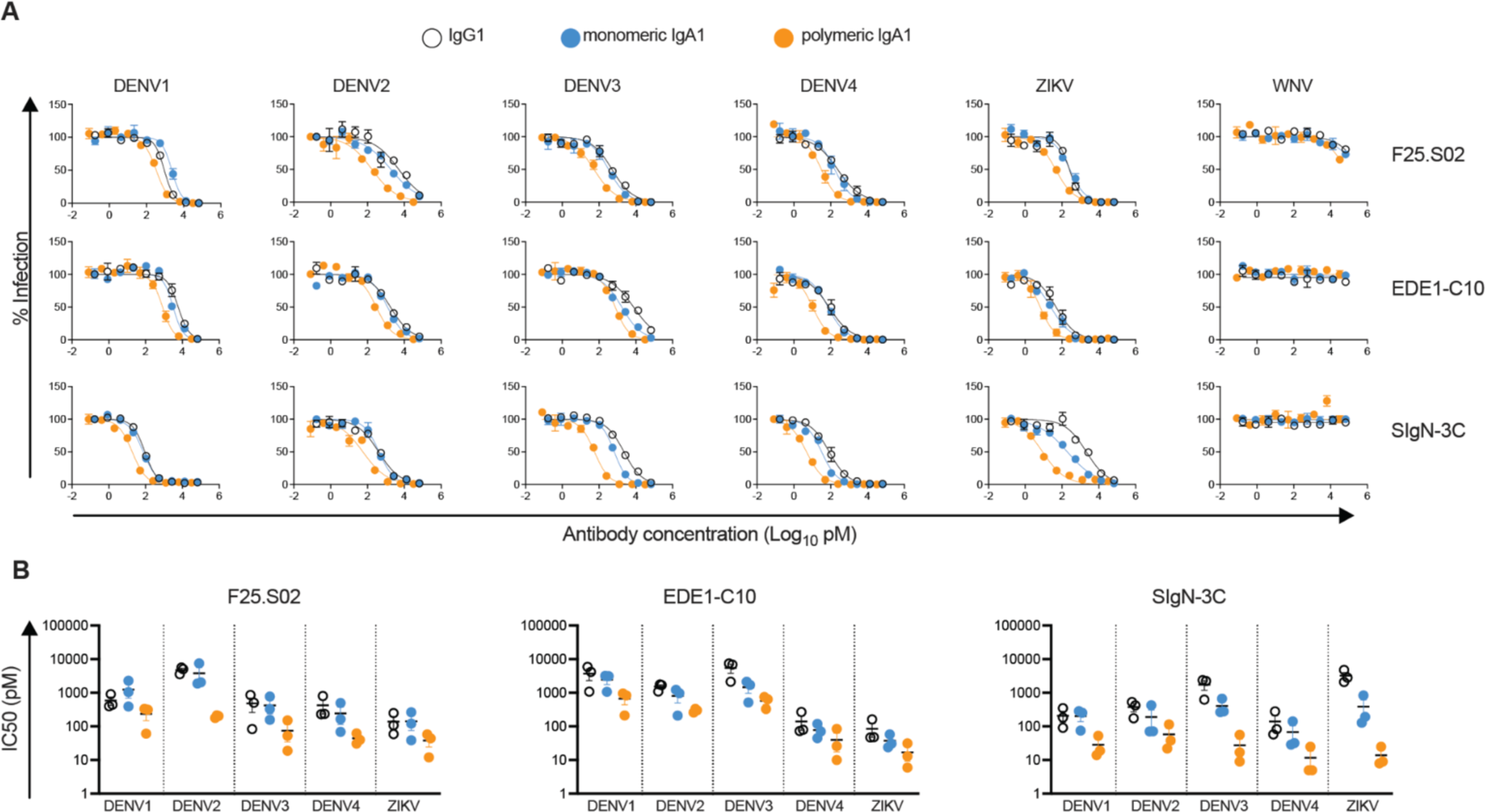
Neutralization profile of antibodies expressed as IgA1. **(A)** IgG1 (open circles), monomeric IgA1 (blue circles), and polymeric IgA1 (orange circles) versions of F25.S02 (top row), EDE1-C10 (middle row), and SIgN-3C (bottom row) were tested for their ability to neutralize reporter virus particles indicated in each column. Dose-response curves are representative of 3 independent experiments, each tested in duplicate wells. Data points and error bars represent the mean and range of the duplicates, respectively. **(B)** Comparison of IC50 values of F25.S02 (left), EDE1-C10 (middle), SIgN-3C (right) expressed as IgG, monomeric IgA1, and polymeric IgA1 against the viruses indicated on the x-axes. Color scheme is similar to (A). Each data point represents an independent experiment in which antibody isotypes were tested in parallel. Horizontal bars indicate the mean. Error bars represent the standard error of the mean.

Antibody expression as polymeric IgA1 further increased potency compared to IgG1 to varying extents. This effect was most apparent for viruses against which the IgG1 version of the particular antibody was the least potent; for F25.S02, EDE1-C10 and SIgN-3C polymeric IgA1, the largest IC50 reduction compared to IgG1 was observed against DENV2 (20-fold), DENV3 (9-fold), and ZIKV (167-fold), respectively (**Figure 6B**). This increased potency of IgA1 bnAbs is unlikely due to non-specific effects as none neutralized the more antigenically distant WNV (**Figure 6A**).

### IgA1 antibodies inhibit enhancement of infection by IgG1

Virtually all IgG antibodies can enhance flavivirus infection *in vitro* at sub-neutralizing concentrations, presumably by facilitating uptake of IgG-virus complexes into FcɣR-expressing cells ^88^. Accordingly, IgG1 versions of newly and previously identified bnAbs enhanced infection to various extents in K562 cells (**Figure S6**) commonly used to study ADE as they express FcɣRIIa (**Figure S7**) and are poorly permissive to flavivirus infection in the absence of IgG ^89^. We did not detect enhancement of ZIKV infection by J9, F09.S05, and F05.S03 (**Figure S7**), suggesting that the lack of ZIKV neutralizing activity by these antibodies (**Figure 3**) could be explained by their inability to bind ZIKV.

Figure 6 demonstrates that regardless of native isotype, F25.S02, EDE1-C10, and SIgN-3C bnAbs expressed as IgA1 retained IgG1 neutralization breadth and potency. As existing studies of ADE of viral infection or disease have focused on the role of IgG-FcɣR interactions ^12,17,21,90–92^, we next investigated the role of IgA in enhancing DENV infection. Specifically, we tested the ability of IgA1 versions of F25.S02, EDE1-C10, and SIgN-3C to enhance DENV1 and DENV4 infection; these viruses were chosen as the infectivity curves obtained across the concentration range of IgG1 versions of bnAbs of interest fully captured both enhancement and neutralization in K562 cells (**Figure S6**).

As expected, IgG1 but not IgA1 versions of F25.S02, EDE1-C10, and SIgN-3C enhanced DENV infection in K562 cells (**Figure 7A**), which do not express Fc alpha receptor (FcɑR1) (**Figure S7**). Surprisingly, monomeric IgA1 antibodies failed to enhance DENV infection even in U937 monocytes (**Figure 7B**), which express FcɑR1 in addition to FcɣRs (**Figure S7**) ^93,94^. Moreover, ADE assays using mixtures of IgG1 and IgA1 antibodies at various ratios demonstrated that autologous IgA1 antibodies inhibited IgG1-mediated ADE of DENV infection in U937 cells (**Figure 7B**) in a dose-dependent manner, as revealed by area under the curve analyses (**Figure 7C**). That this effect was observed for all 3 bnAbs regardless of native isotype and epitope specificity indicates that IgA1 antibodies can broadly interfere with IgG1-mediated ADE. Crucially, an isotype control IgA1 antibody had virtually no effect on ADE mediated by IgG1, indicating that inhibition was due neither to a reduction in IgG1 concentration in IgG1/IgA1 mixtures nor the presence of non-specific IgA1. Rather, IgA1 inhibits ADE mediated by IgG1 likely via direct competition of binding to virions.

**Figure 7.**
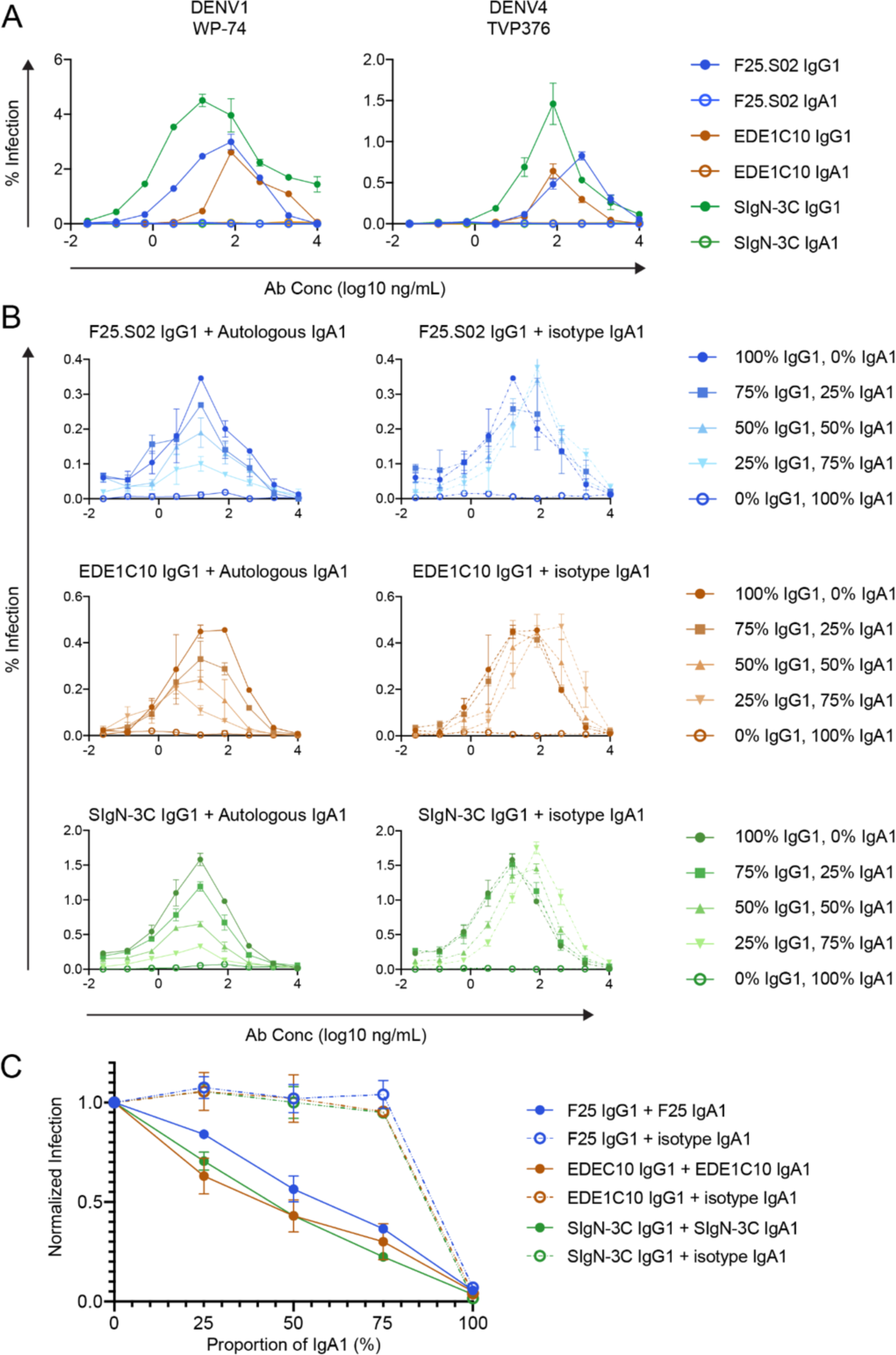
Effect of antibody isotype on antibody dependent enhancement (ADE). In (A-B), DENV1 (left panel) and DENV4 (right panel) reporter virus particles were pre-incubated with serial dilutions of IgG1 and/or IgA1 forms of the indicated antibodies prior to infection of target cells expressing Fc receptor for IgG and/or IgA. **(A)** Dose-response ADE assays in K562 cells, which express FcγRII but not FcαRI. IgG1 and IgA1 antibodies were tested individually. The data points and error bars represent the means and range of duplicate infection, respectively. **(B)** Dose-response ADE assays in U937 monocytes, which express both FcγRII and FcαR. F25.S02 (top row), EDE1-C10 (middle row) or SIgN-3C (bottom row) IgG1 was mixed with autologous IgA1 or an IgA1 isotype control at the indicated ratios by mass before serial dilution and pre-incubation with virus. The experiment was performed twice in duplicate wells and a representative experiment is shown. The data points represent the means of the duplicates and the error bars the range. **(C)** Area under the curve analysis for experiments represented in (B). For both experimental replicates the area of the curve for each infection condition was calculated and normalized to infection in the 100% IgG1 condition.

## DISCUSSION

Unlike most antibody discovery approaches that involve screening large panels of antibodies expressed by sorted and baited B cells ^38^, we previously established a proof-of-concept for a bioinformatics-based strategy to identify not only antigen-specific antibodies, as shown previously by other groups ^50,95,96^, but also those with broadly neutralizing activity ^39^. Here, we have improved upon our previous approach and leveraged scRNAseq of B cells to identify multiple antibodies that broadly and potently neutralized DENV1-4 and in some cases, ZIKV. Previous studies characterizing flavivirus bnAb responses have used antibody isolation protocols that specifically enriched the IgG isotype ^28,33–35^. In contrast, our scRNAseq approach is designed to capture full-length antibody sequences in an unbiased manner. Although most new bnAbs we discovered were of the IgG1 isotype, consistent with previous findings ^27,28,39^, we also describe for the first time an IgA1 antibody with broadly neutralizing activity against DENV1-4 and ZIKV.

Despite broad and potent serum neutralizing activity in all 4 donors selected for antibody repertoire analysis, almost all monoclonal bnAbs were isolated from only one donor (014). Although we did not set out to formally investigate the basis for donor-dependent effects, consistent with previous findings ^45,48^, antibody neutralizing activity could be partly explained by sample collection time (**Figure S1**), which likely affected our ability to capture transiently circulating plasmablasts (**Figure 2A**), many of which encode bnAbs ^25,27,28,39^. Alternatively, the observed serum neutralization breadth and potency across donors was due to a combination of antibodies with multiple specificities. However, within a given donor, we did not detect an obvious pattern of complementary neutralizing activity among antibodies from distinct clonal families to support this hypothesis (**Figure S2**). The number and order of prior flavivirus exposures also impact bnAb development ^21^. It is interesting that unlike other donors analyzed, donor 014 was confirmed to have been acutely co-infected with two DENV serotypes (**Figure S1**). Prior studies have documented concurrent infection by multiple DENV serotypes in hyperendemic regions ^97–102^, but whether co-infection uniquely impacts bnAb induction has not been systematically explored. Finally, while we successfully identified multiple new bnAbs, our *in silico* down-selection criteria are likely subject to stochastic processes to some degree ^103^.

Although neutralizing activity is thought to be primarily determined by somatic hypermutation within IgG antibody variable regions, Fc isotype can also impact neutralization potency and/or breadth against many viruses ^82–86^. For example, a recent study described a naturally occurring ZIKV-specific pentameric IgM antibody (DH1017.IgM) whose potency depended on the IgM isotype ^86^. Unlike DH1017.IgM, which did not neutralize DENV, here we identified F25.S02, an IgA1 antibody that potently cross-neutralized ZIKV and DENV1-4 and retained its potency as IgG1. While IgA bnAbs have been described for other antigenically distinct viruses such as HIV ^81^ and SARS-CoV-2 ^83^, to our knowledge, F25.S02 is the first known IgA bnAb against flaviviruses. In addition to its distinct isotype, our epitope mapping results demonstrate that despite some similarities, F25.S02 has unique binding and neutralization determinants compared to EDE1-C10 ^28,30,31^ and SIgN-3C ^27,36,37^ IgG1 antibodies, which represent the only 2 known classes of bnAbs that potently neutralize ZIKV and DENV1-4.

Human IgA antibodies in serum and mucosal sites exist primarily as monomeric or dimeric/polymeric forms, respectively ^87^. As monomeric IgA1, F25.S02 displayed comparable neutralizing activity to IgG1 against DENV1-4 and ZIKV. In contrast, we show that expression of EDE1-C10 and SIgN-3C bnAbs as monomeric IgA1 improved potency against some viruses, despite their native IgG1 isotype ^28,40^. These findings are consistent with epitope- and virus-dependent effects of antibody isotype on neutralization ^81^. Expression of all 3 bnAbs as polymeric IgA1 increased potency against DENV1-4 and ZIKV relative to corresponding monomeric IgA1 or IgG1 versions. Defining the mechanism(s) behind this observation awaits further studies but it suggests that the epitope arrangement of these bnAbs allows multivalent engagement by polymeric IgA on the same virion. Alternatively or in addition to this mechanism, polymeric IgA could bind the same epitope on multiple virions to cause aggregation. Both mechanisms of virion engagement have been shown for DH1017.IgM, depending on the particular antibody conformation ^86^.

Compared to other isotypes, IgA1 antibodies have a greater distance between Fabs relative to each other and to the Fc domain ^104,105^, providing a possible mechanism for unique neutralizing and Fc-dependent effector functions^81^. Further, engagement of IgA with FcɑR1 is distinguished from that of other isotypes with their Fc receptors in terms of stoichiometry, orientation, and protein binding sites ^106^, which could impact the efficiency with which different antibody isotypes facilitate ADE. Consistent with this hypothesis, we demonstrate that unlike IgG1, IgA1 versions of bnAbs failed to mediate ADE of DENV infection even in cells expressing Fc receptors for both isotypes. Moreover, IgA1 antibodies not only displayed neutralization breadth and potency comparable or superior to IgG1, but also inhibited IgG1-mediated ADE in a dose-dependent manner, likely via competition for binding to virions. Thus, we extend previous results demonstrating the ability of monomeric IgA1 to antagonize IgG-mediated ADE of DENV in cells that express FcɣR but not FcɑR1 ^107^.

Existing studies of flavivirus immunity have heavily focused on the role of IgG antibodies and their interactions with FcɣRs ^12,16,17,21,90,108^. Although the *in vivo* relevance of our results remains to be validated, they nevertheless highlight an underappreciated role for flavivirus-specific IgA antibodies in infection and immunity. Indeed, recent studies reported a high proportion of DENV-reactive IgA-expressing plasmablasts following acute primary infection and to a lesser extent, secondary infection ^48,109^. Our analysis of circulating B cell repertoires here also demonstrates that while IgG dominated the response, IgA and IgM antibodies were prevalent (**Figure 2B**). Notably, FcɑR1 is expressed on myeloid cells, including monocytes, macrophages, and dendritic cell subsets ^110–113^, all of which also express FcɣRs and are thought to be principal target cells for DENV *in vivo* ^41,114–118^. Intriguingly, IgA-FcɑR1 interactions can modulate activating or inhibitory responses mediated by *other* Fc receptors ^119,120^. Together, these observations underscore the importance of future studies to account for the complex interplay among distinct antibody isotypes and Fc receptors in modulating flavivirus immunity and pathogenesis. Determining whether IgA and other non-IgG isotypes mitigate or potentiate antibody-associated disease *in vivo* will inform strategies to improve the safety and efficacy of antibody-based countermeasures ^121^.

A limitation of our study is that we did not evaluate the *in vivo* protective and pathogenic potential of identified bnAbs, in part due to the lack of an animal model that fully recapitulates dengue immunity and disease ^122–124^. Evaluating these properties for IgA antibodies in existing mouse models is especially challenging as they do not express FcɑR1 homologs ^125^. Thus, cohort studies similar to those that have defined IgG-associated correlates of protection or disease ^12,13,17,21^ would be most informative. Another limitation is that we analyzed antibody repertoires from a relatively small donor sample size. Additionally, because our primary goal was to discover bnAbs, we focused on antibodies encoded by transiently circulating plasmablasts, which often display neutralization breadth and potency. Although there is functional overlap between the DENV-specific plasmablast antibody repertoire with that of memory B cell and long-lived plasma cell subsets ^46^, future studies will need to determine whether the bnAbs we identified here contribute to durable immunity.

## METHODS

### Cohort Samples

The study’s use of samples from DENV and ZIKV infected human donors was approved by the Stanford University Administrative Panel on Human Subjects in Medical Research (Protocol #35460) and the Fundación Valle del Lili Ethics committee in biomedical research (Cali, Colombia). All participants, their parents, or legal guardians provided written informed consent, and subjects 6 years of age and older provided assent. We collected blood samples from individuals who presented with symptoms compatible with dengue between 2016 and 2017 to the Fundación Valle del Lili in Cali, Colombia. Each blood sample was centrifuged to separate serum and peripheral blood mononuclear cells (PBMCs). Sera was stored at −80°C and corresponding PBMCs were cryopreserved and stored in liquid nitrogen. Cohort details have been previously described ^41,42^.

### Cell lines

Expi-CHO-S Cells (Cat# A29127; ThermoFisher Scientific, Waltham MA) were cultured in ExpiCHO Expression Medium (Cat# A2910001; ThermoFisher Scientific) and maintained at 37°C in 8% CO2 on a platform rotating at 125 rpm with a rotational diameter of 19 cm. They were subcultured according to the manufacturer’s instructions. HEK-293T/17 cells (Cat# CRL-11268, ATCC, Manassas, VA) and Vero-C1008 cells (Cat# CRL-1586, ATCC) were maintained in DMEM (Cat# 11965118; ThermoFisher Scientific) supplemented with 7% fetal bovine serum (FBS)(Cat# 26140079, lot 2358194RP, ThermoFisher Scientific) and 100 U/mL penicillin-streptomycin (Cat# 15140–122; ThermoFisher Scientific). Raji cells stably expressing DCSIGNR (Raji-DCSIGNR) ^126^ (provided by Ted Pierson, NIH), K562 cells (Cat# CCL-243, ATCC), and U937 cells (Cat# CRL-1593.2, ATCC) were maintained in RPMI 1640 supplemented with GlutaMAX (Cat# 72400–047; ThermoFisher Scientific), 7% FBS and 100 U/mL penicillin-streptomycin. C6/36 cells (Cat# CRL-1660, ATCC) were maintained in EMEM (Cat# 30–2003, ATCC) supplemented with 10% FBS at 30 °C in 5% CO2. All cell lines were maintained at 37 °C in 5% CO2 unless otherwise stated.

### Preparation of cells for single-cell RNA sequencing

Cryopreserved PBMCs were thawed quickly in a 37°C water bath and transferred to a 50 mL conical tube. Thirty mL of RPMI 1640 supplemented with 10% FBS (no antibiotics) was added to the cells dropwise while gently swirling. Cells were counted and CD19+ B cells were isolated using the EasySep Human Pan-B cell enrichment kit (Cat# 19554, StemCell Technologies, Vancouver, Canada) according to the manufacturer’s instructions. The resulting cells were incubated in a cocktail containing a live/dead stain (Cat# L34957, Thermo Scientific) and fluorescently labeled antibodies for CD20-eFluor450 (Cat# 48-0209-42, Invitrogen, Waltham, MA), CD38-FITC (Cat# 303504, Biolegend, San Diego, CA), CD27-PE-Cy7 (Cat# 25-0271-82, Invitrogen), CD19-APC (Cat# 555415, BD Biosciences, Franklin Lakes, NJ), CD3-APC-Cy7 (Cat# 300318, Biolegend), CD8-APC-Cy7 (Cat# 344714) and CD14-APC-Cy7 (Cat# 301820) for 30 min at 4°C. Stained cells were washed twice in FACS wash buffer (10% FBS in PBS) and strained through FACS tubes with strainer caps (Cat# 352235, BD Biosciences). The cells were analyzed on a BD FACS Aria flow cytometer to assess the purity of B cells (CD19+) and determine the fraction of cells that were plasmablasts (CD3^-^, CD8^-^, CD14^-^, CD19 ^mid^ ^to^ ^hi^, CD20^-^, CD27^+^, CD38^+^). If the fraction of plasmablasts in the B cell sample was <10% (Donor 012) we sorted plasmablasts via flow cytometry. If the fraction of plasmablast in the B cell sample was >10% (Donors 001, 002, 014), we proceeded without further enrichment.

The cells were prepared for RNA library generation using the Chromium Next GEM Single Cell 5’ Library and Gel Bead Kit v1.1 (Cat# PN-1000167, 10X Genomics, Pleasanton, CA) according to the manufacturer’s instructions. A library enriched for variable regions of B cell receptors (BCR library) was generated using the Chromium Single Cell V(D)J Enrichment Kit, Human B Cell (Cat# PN-1000016, 10X Genomics) and the global gene expression library (GEX library) was generated using the Chromium Single Cell 5’ Library Construction Kit (Cat# PN-1000020, 10X Genomics), both according to the manufacturer’s instructions. Both libraries from the sample D014 were sequenced on an Illumina HiSeq, The libraries for the samples D001 (donor 001), D002 (donor 002), and D012 (donor 012) were sequenced on an Illumina NovaSeq 6000. Sequencing data were demultiplexed and aligned to the human transcriptome GRCh38-2020-A using cellranger (10X genomics) version 5.0.1 (D001, D002, D012) or 5.0.0 (D014, donor 014), which also identified the isotype of each antibody. The “filtered” cellranger output was then passed to *partis* for paired heavy/light chain clustering and annotation with default parameters ^51^. This included the default *partis* disambiguation of incomplete and ambiguous heavy/light pairing information, which for instance resolved an atypically large number of droplets in D014 with reads from more than one cell. After grouping all sequences from an individual donor into clonal families, *partis* estimated the V, D, and J gene segments that composed the naive antibody sequence. B cell subtypes were identified using previously described gene markers ^48^ in the AUCell package (1.12.0). Isotype annotations were taken from the cellranger output.

### Selection of candidate bnAbs from single-cell RNA sequencing data

The variable regions of the paired heavy and light chain sequences were grouped into clusters based on inferred shared ancestry (clonal families) using *partis*, as described previously ^51^. For the first round of screening intended to find families that encode bnAbs, we selected the largest clonal families from each donor excluding those in which the mean somatic hypermutation (measured by nucleotide sequence) was below 2%. Within the selected families we selected 1-2 sequences that had the lowest Hamming distance to consensus (i.e. the sequence consisting of the most common amino acid present at each position), excluding those that were not encoded by plasmablasts. The selected antibodies were screened for their ability to neutralize DENV1-4 and ZIKV (described below) and those that neutralized >50% of infection of 3 or more viruses were considered “hits”. We initiated a second round of screening of antibodies from clonal families that had produced hits in the first round. Within each family we selected antibodies in ascending order of Hamming distance to the consensus, again excluding those that were not encoded by plasmablasts.

### Expression of recombinant antibodies

Heavy and light chain constructs for recombinant MZ4 IgG1 expression have been described previously ^33^ and were provided by Shelly Krebs (Walter Reed Army Institute of Research). For other antibodies, heavy and light chain variable regions were synthesized (Twist Bioscience, South San Francisco, CA). Variable region sequences for newly identified antibodies were selected from our scRNAseq data; those for control bnAbs were determined based on the protein database (PDB) entries 4UT9 (EDE1-C10), 4UTA (EDE1-C8), 4UT6 (EDE2-B7), 4UTB (EDE2-A11), and 7BUD (SIgN-3C). All variable regions were cloned into the expression vectors provided by Patrick Wilson (University of Chicago): AbVec-hIgG1 (GenBank accession # FJ475055), AbVec-hIgKappa (GenBank accession# FJ475056) and AbVec-hIgLambda (GenBank accession # FJ517647), respectively. The variable regions were synthesized with overlapping sequences to their respective vectors. The sequence that was appended to the 5’ end was the same for all vectors: TAGTAGGAACTGCAACCGGTT. The sequence appended to 3’ ends was specific to each vector: for AbVec heavy: CGGTCGACCAAGGGCCCATCGG, for AbVec kappa: CGTACGGTGGCTGCACCATC, and for AbVec lambda: GGTCAGCCCAAGGCCAACCCCACTGTCACTCTGTTCCCACCCTCGAGTGAGGAGCTTC AAGC. Heavy, kappa, and lambda vectors were linearized by digestion with SalI/AgeI, BsiWI/AgeI, and XhoI/AgeI, respectively as described ^127^. Synthesized fragments and linearized vectors were ligated using NEBuilder HiFi DNA Assembly Master Mix (Cat# E2612L, New England Biolabs, Ipswich, MA) according to the manufacturer’s instructions.

IgA1 heavy chains were generated by cloning the variable regions of selected antibodies into the expression vector pFUSEss-CHIg-hA1 (Cat# pfusess-hcha1, Invivogen, San Diego, CA). Variable regions of the antibody coding sequences were PCR amplified using the IgG1 heavy chain expression plasmid as a template and custom primers that appended an EcoRI site and an NheI site at the 5’ and 3’ ends respectively. Primer sequences were as follows: for F25.S02 GTACACGAATTCGCAGGTGCAGCTGGTGC (forward) and GACTCTGCTAGCTGAGGAGACGGTGACC (reverse); for EDE1-C10: GTACACGAATTCGGAGGTCCAACTTGTTG (forward) and GACTCTGCTAGCA GAGCTTACGGTTACG (reverse); and for SIgN-3C GTACACGAATTCGGAAGTACAACTGGTGC (forward) and GACTCTGCTAGCTGAACTAACAGTTACCAG (reverse). The PCR amplicons and the vector were digested with EcoRI and NheI and the resulting fragments were ligated using T7 DNA ligase (Cat# M0318, New England Biolabs).

All AbVec antibody expression plasmids (IgG1-heavy, kappa, and lambda) were confirmed by Sanger sequencing using the primer “AbVec sense”: GCTTCGTTAGAACGCGGCTAC. IgA1 expression plasmids were confirmed by whole plasmid nanopore sequencing (Plasmidsaurus, Eugene, OR). To produce IgG1 and monomeric IgA1, heavy and light chain expression vectors were co-transfected into cultures of ExpiCHO-S cells at 0.8 ng/mL total DNA concentration at 1:1 mass ratio using OptiPro serum free medium (Cat#12309, Gibco) and Expifectamine CHO Transfection Kit (Cat# A29130, Gibco) according to the manufacturer’s instructions. To produce IgA1 dimers, plasmids encoding heavy, light, and joining chain (Cat# pUNO4-hJCHAIN, InvivoGen) were co-transfected at 0.8 ng/mL total DNA concentration at 1:1:1 mass ratio using the same medium and transfection reagents. Supernatant containing secreted antibodies was collected 8 days post transfection, centrifuged at 3220 x g for 10 minutes and filtered through a 0.45 µm Steriflip filter (Cat# SE1M003M00, Millipore-Sigma).

### Purification of antibodies

The hybridoma D1-4G2-4-15, which expresses the antibody 4G2 was obtained from ATCC (Cat# HB-112). The hybridoma was expanded and IgG was purified from culture supernatant by the Fred Hutchinson Cancer Center Antibody Technology Core. The purified antibody was conjugated to APC using the Lighting-Link APC-conjugation kit (Cat# ab201807, Abcam) according to the manufacturer’s instructions. Recombinant IgG1 produced in transfected ExpiCHO-S cells as described above was purified using MabSelect Sure LX protein A agarose beads (Cat# 17-5474-01, Cytiva Life Sciences, Marlborough, MA) according to the manufacturer’s instructions. Recombinant IgA1 produced in ExpiCHO-S cells as described above was purified using protein M agarose beads (Cat# gel-pdm-2, InvivoGen US) according to the manufacturer’s instructions. IgA1 multimers were separated from monomers via size exclusion chromatography on a HiLoad 16/600 Superdex 200 pg column using 70 mL PBS as the eluate. A monomeric IgA1 antibody (Cat# 31148, ThermoFisher) was used as a standard for SDS-PAGE and as a negative control for ADE assays as indicated.

### Production of reporter virus particles

Reporter virus particles were produced by co-transfection of HEK-293T/17 cells with (i) a plasmid expressing a WNV subgenomic replicon encoding GFP in place of structural genes ^128^, and (ii) a plasmid encoding C-prM-E structural genes from the following viruses: DENV1 Western Pacific-74 (WP-74) ^129^, DENV1 16007 ^130^, DENV2 16681 ^129^, DKE-121 ^70^, WNV NY99 ^128^, and ZIKV H/PF/2013 ^131^. Briefly, 8 x 10^5 HEK-293T/17 cells were plated in each well of a 6-well plate, The following day each well was co-transfected with 1 µg of replicon-encoding plasmid and 3 µg of C-prM-E-encoding plasmid using Lipofectamine 3000 (Cat# L3000-015; ThermoFisher Scientific) according to the manufacturer’s instructions. Four hours post-transfection, media was replaced with low-glucose DMEM (Cat# 12320–032; ThermoFisher Scientific) containing 7% FBS and 100 U/mL penicillin-streptomycin (i.e. low-glucose DMEM complete) and cells were transferred to 30 °C in 5% CO2. Virus-containing supernatant was harvested twice per day at days 3 through 8 post-transfection, centrifuged at 700 x g for 5 min. The clarified supernatant was passed through a 0.45 µm Steriflip filter (Cat# SE1M003M00, Millipore-Sigma, St. Louis, MO), pooled, aliquoted, and stored at −80 °C.

Reporter virus particles with increased efficiency of prM cleavage were produced as above by co-transfecting plasmids encoding the replicon, structural genes, and human furin (provided by Ted Pierson, NIH) at a 1:3:1 mass ratio. DENV3 strain CH53489 (Cat# RVP-301; Integral Molecular, Philadelphia, PA) and DENV4 strain TVP376 reporter viruses (Cat# RVP-401; Integral Molecular) were obtained commercially and were produced by co-transfection of the DENV3 or DENV4 CprME plasmid with the DENV2 strain 16681 replicon as previously described ^132^.

Infectious titers of reporter viruses were determined by infection of Raji-DCSIGNR cells. At 2 days post-infection, cells were fixed in 2% paraformaldehyde (Cat# 15714S; Electron Microscopy Sciences, Hatfield, PA), and GFP positive cells quantified by flow cytometry (Intellicyt iQue Screener PLUS, Sartorius AG, Gottingen, Germany).

### Generation of E protein variants

Construction of DENV2 16681 reporter virus variants in which E protein sites were substituted with corresponding ZIKV H/PF/2013 amino acid residues individually or in combination have been previously described. Here, we used similar methods to generate individual alanine mutations. Specifically, the DENV2 16681 CprME expression construct ^129^ was used as a template for Q5 site-directed mutagenesis (Cat# E0554S; New England Biolabs, Ipswich, MA) and primers generated by NEBaseChanger (New England Biolabs, Ipswich, MA). The entire plasmid was sequenced (Plasmidsaurus, Eugene, OR) to confirm the presence of the desired mutation(s) only.

### ELISA

DENV2 16681 reporter virus particles were concentrated by ultracentrifugation through 20% sucrose at 166,880 x g for 4 hr at 4 °C, resuspended in 1/100 volume of HNE buffer (5 mM HEPES, 150 mM NaCl, 0.1 mM EDTA, pH 7.4), and stored at −80 °C. Nunc 384-Well Clear Polystyrene Plates (Cat# 164688 ThermoFisher) were coated with 20 uL/well of recombinant E monomers (Cat#DENV2-ENV, Native Antigen Co, Kidlington, United Kingdom) at 3 µg/mL or 20 µL/well of antibody 4G2 at 50 ug/mL overnight. The next day plates were washed once with 50 µL wash buffer (0.05% Tween-20 in PBS) and blocked with 50uL of blocking buffer (3% nonfat milk in PBS) at 37°C for 45 min. Blocking buffer was aspirated from wells that had received 4G2 and replaced with 20uL of 100X concentrated reporter virus particles diluted 1:1 in blocking buffer. Wells that had received E monomers were left in blocking buffer and plates were incubated at 37°C for 45 min. Wells were washed 3 times with 50 uL of wash buffer, received 30 µL of primary antibody at 100 µg/mL, and were incubated at 37°C for 45 min. Wells were washed 6 times with 50 µL wash buffer, received 30 µL of mouse anti-human Ab (Cat# 05-4220, ThermoFisher) at 1 µg/mL, and were incubated at 37*C for 45 min. Finally, wells were washed 6 times with 50 µL wash buffer, received 30 µL of TMB (Cat# 34028 ThermoFisher), and were incubated until a color change was apparent. The reaction was stopped with 15 uL of 1N HCl and absorbance at 450 nm was read on SpectraMax i3x plate reader (Molecular Devices, San Jose, CA)

### Binding screen against alanine library

We screened binding of antibodies F25.S02 and F05.S03 to a DENV2 16681 library where each prM/E polyprotein residue was mutated to alanine (or alanine residues to serine) ^79^. In total, 559 sequence confirmed DENV2 mutants (99.6% coverage of the prM/E protein) were arrayed into 384-well plates (one mutation per well). The optimal screening condition was determined using an independent immunofluorescence titration curve against wild-type prM/E expressed in HEK293T cells to ensure that signals were within the linear range of detection and that signal exceeded background by at least 5-fold. F25.S02 and F05.S03 bound sufficiently well for screening only when the prM/E expression plasmid was co-transfected with a furin expression plasmid to enhance cleavage of prM to M. Thus, for antibody screening, plasmids encoding the DENV protein variants were individually co-transfected with furin expression plasmid into HEK-293T cells and expressed for 22 hr before incubation with purified IgG1 antibodies (0.1-2.0 µg/mL) diluted in 10% normal goat serum (NGS) (Sigma-Aldrich, St. Louis, MO) in PBS plus calcium and magnesium (PBS++).

Antibodies were detected using 3.75 µg/mL Alexa Fluor 488-conjugated secondary antibody (Jackson ImmunoResearch Laboratories) in 10% NGS. Cells were washed three times with PBS++ followed by 2 washes in PBS, then fixed in 4% paraformaldehyde, washed in PBS (Electron Microscopy Sciences), and resuspended in Cellstripper (Cat# 25-056-CI, Corning Inc, Corning, NY) plus 0.1% BSA (Sigma-Aldrich). Mean cellular fluorescence was detected by flow cytometry (Intellicyt iQue Screener PLUS, Sartorius AG).

Antibody reactivity against each mutant was calculated relative to reactivity with wild-type prM/E, by subtracting the signal from mock-transfected controls and normalizing to the signal from wild-type protein-transfected controls. The entire library data for each antibody was compared to control antibodies. Mutations were identified as critical to the antibody epitope if they did not support reactivity of the test antibody, but supported reactivity of other control antibodies. This counter-screen strategy facilitates the exclusion of DENV prM/E protein mutants that impact folding or expression.

### Neutralization and antibody-dependent enhancement assays using reporter virus particles

All neutralization and ADE assays using the following strains were performed with reporter virus particles: DENV1 West-Pac 74, DENV1 16607, DENV2 16681, DENV3 CH53489, DENV4 TVP376, DKE-121, ZIKV H/PF/2013. Depending on the assay, stocks of reporter virus particles diluted to 5–10% final infectivity were incubated with either heat-inactivated serum (56 °C for 30 min), 1/10 diluted ExpiCHO-S cell supernatant containing recombinant IgG1, or 5-fold serial dilutions of purified monoclonal antibodies for 1 hr at room temperature before addition of 2 x 10^5 Raji-DCSIGNR cells (neutralization assays), K562 cells (ADE assays), or U937 cells (ADE assays). After 48 hr incubation at 37 °C, cells were fixed in 2% paraformaldehyde and GFP positive cells were quantified by flow cytometry (Intellicyt iQue Screener Plus, Sartorius AG). For experiments using single dilutions of serum or ExpiCHO-S cell supernatant, infection was normalized to conditions without serum/supernatant and expressed as % infection of the untreated condition. For experiments using serial dilutions of serum or of purified monoclonal antibodies, infection was normalized to conditions without serum/antibody and analyzed by non-linear regression with a variable slope and the bottom and top of the curves constrained to 0% and 100%, respectively (Graph-PadPrism v8, GraphPad Software Inc). Results from experiments using serially diluted serum were reported as the reciprocal dilution at which 50% of infection was neutralized (NT50). Results from experiments using serially diluted purified antibodies were reported as the concentration at which 50% of infection was neutralized (IC50).

### Production, titer and neutralization of fully infectious virus

DENV1 UIS 998 (isolated in 2007, Cat# NR-49713), DENV2 US/BID-V594/2006 (isolated in 2006, Cat# NR-43280), DENV3/US/BID-V1043/2006 (isolated in 2006, Cat# NR-43282), DENV4 strain UIS497 (isolated in 2004, Cat# NR-49724) were obtained from BEI Resources (Manassas, VA). Viral stocks were expanded by infecting 70% confluent C6/36 cells and virus-containing supernatant was collected and pooled at days 3 to 8 post infection. DENV4 H241 (isolated 1956, Cat# TVP17463) was obtained from the World Reference Center for Emerging Viruses and Arboviruses at the University of Texas Medical Branch (Galveston, TX). The seed stock was expanded by infecting 90% confluent Vero cells and virus-containing supernatant was collected 7 days post infection. All virus-containing supernatants were centrifuged at 500 x g for 5 min, filtered through a 0.45 µm Steriflip filter (Cat# SE1M003M00, Millipore-Sigma), and stored at −80°C. Viral stocks were titered by infecting 2e5 Raji-DCSIGNR cells with 2-fold serial dilutions. Two days post infection cells were fixed and permeabilized using BD cytofix/cytoperm (Cat# 554717, BD Biosciences) according to the manufacturer’s instructions before being incubated with APC-conjugated 4G2 for 30 minutes at 4°C. Cells were washed twice in cytoperm/wash buffer and APC+ positive cells were quantified by flow cytometry.

For dose response neutralization assays using fully infectious virus, stocks were diluted to achieve 5-10% infection in Raji-DCSIGNR cells were incubated with 5-fold serial dilutions of antibodies for 1 hour, then combined with 2e5 Raji-DCSIGNR cells and incubated at 37°C 5% CO2, before being stained for E protein as described above. IC50 values were calculated as described above for neutralization assays using reporter virus particles.

### Determining Fc receptor expressio

K562 cells and U937 cells were washed in FACS wash (FW, 2% FBS in PBS) and resuspended in 50 µL of staining or isotype control antibody and incubated at 4°C for 30 min. For FcγRII we stained with anti-CD32-FITC (Cat# 60012.FI, StemCell) and corresponding mouse IgG2b-FITC isotype control (Cat# 11-4732-81, ThermoFisher Scientific). For FcαRI we stained with anti-CD89/-PE (cat# 555686, BD Biosciences) and corresponding mouse mouse IgG1-PE isotype control (cat# 12-4714-42, ThermoFisher Scientific). Cells were washed twice in FW and analyzed by flow cytometry.

### Statistical analysis

All data were analyzed and plotted in Prism 8.4.3 (GraphPad Software, San Diego, CA).

## ACKNOWLEDGEMENTS

We thank cohort participants and staff; John McNevin, Andrew Berger, and Brian Raden, for assistance with cell sorting; Adam Waickman for advice on scRNAseq analysis; Patrick Wilson for providing IgG1 expression vectors, Ted Pierson for providing Raji-DCSIGNR cells and constructs for reporter virus production; Leah Homad and Andrew McGuire for assistance with polymeric IgA1 purification; Michael Diamond for providing E60 and ZV-67 antibodies; Shelly Krebs for providing expression constructs for antibody MZ4; and Dennis Burton for providing antibody PGT121.

Fully infectious DENV1-4 isolates were obtained through BEI Resources, NIAID, NIH, as part of the World Reference Center for Emerging Viruses and Arboviruses program (WRCEVA).

Molecular graphics and analyses of DENV2 E dimer were performed with UCSF ChimeraX, developed by the Resource for Biocomputing, Visualization, and Informatics at the University of California, San Francisco, with support from National Institutes of Health R01-GM129325 and the Office of Cyber Infrastructure and Computational Biology, National Institute of Allergy and Infectious Diseases.

This work was supported by the Fred Hutchinson Cancer Center Translational Data Science Integrated Research Center New Collaborations Award (LG, FAM, JL, LML, DKR); NIH R01 AI146028 (DKR, FAM); the Howard Hughes Medical Institute (FAM); Viral Pathogenesis and Evolution Training Grant T32 AI083203 (LB); Fred Hutchinson Cancer Center Diverse Trainee Fund (MC); an Investigator Initiated Award W81XWH1910235 from the Department of Defense Office of the Congressionally Directed Medical Research Programs (SE); Catalyst and Transformational Awards from Dr. Ralph & Marian Falk Medical Research Trust (SE); NIH U19 AI057229 supplement (SE); the Chan Zuckerberg Biohub (SE); the Antibody Technology (RRID:SCR_022608), Flow Cytometry (RRID:SCR_022613), and the Genomics & Bioinformatics (RRID:SCR_022606) Shared Resource Facilities of the Fred Hutch/University of Washington/Seattle Children’s Cancer Consortium (P30 CA015704); and the Scientific Computing Infrastructure at Fred Hutch (ORIP grant S10OD028685).

FAM is an Investigator of the Howard Hughes Medical Institute. SE is a Chan Zuckerberg Biohub - San Francisco Investigator. VD was supported by a Chan Zuckerberg Biohub Collaborative Postdoctoral Fellowship.

## SUPPLEMENTAL INFORMATION

**Table S1.**
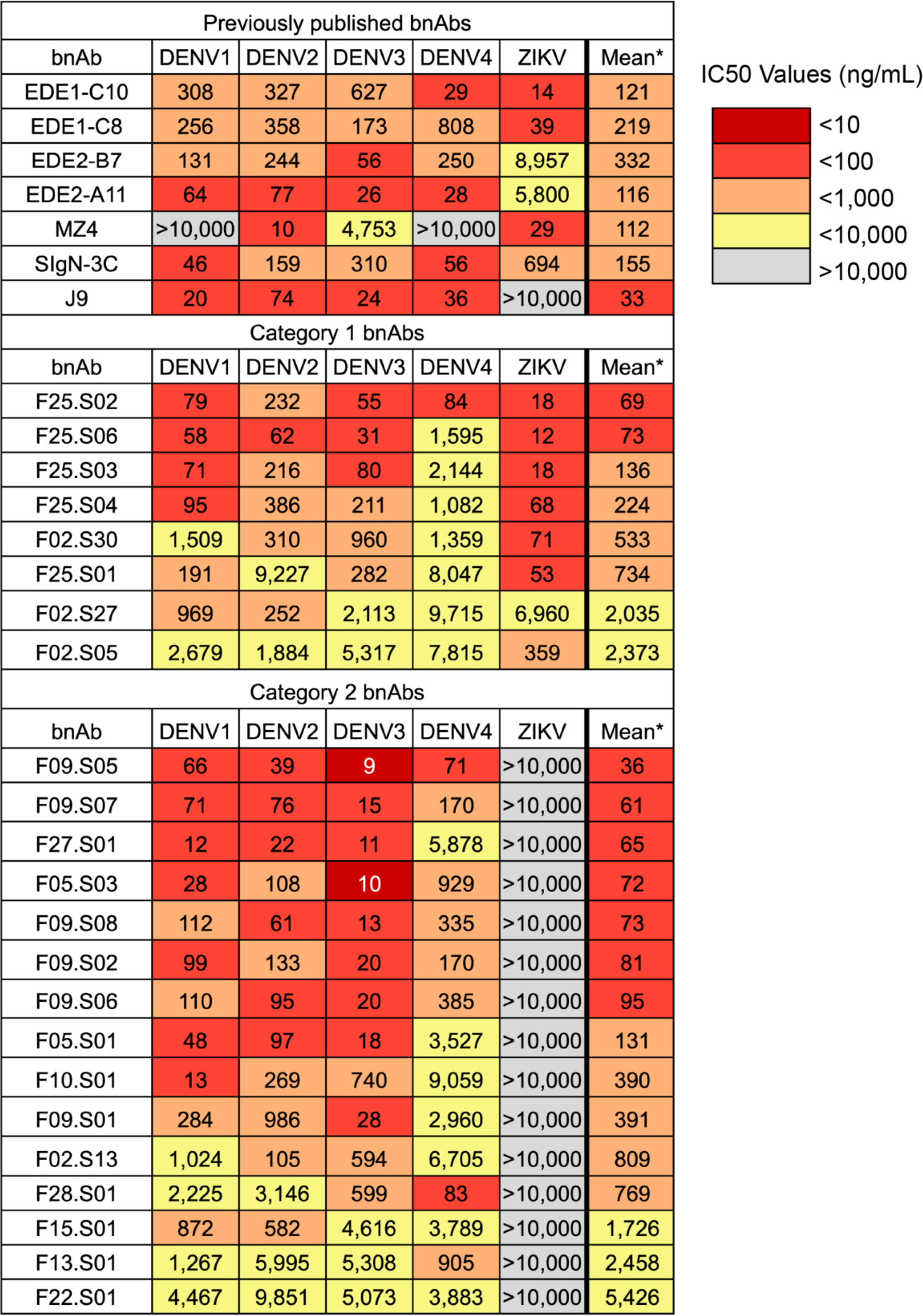
Heatmaps of IC50 values obtained from dose-response neutralization assays using previously published bnAbs, novel category 1 bnAbs, which neutralize DENV1-4 and ZIKV, and novel category 2 novel bnAbs, which neutralize DENV1-4 but not ZIKV. For each virus, the value reported is the arithmetic mean IC50 from at least three independent experiments performed in duplicate. *Geometric mean IC50 for all neutralized viruses, i.e. values >10,000 ng/ml (the highest antibody concentration tested) were omitted. All antibodies were isolated from donor 014 except for F15.S02, which was isolated from donor 012.

**Table S2.**
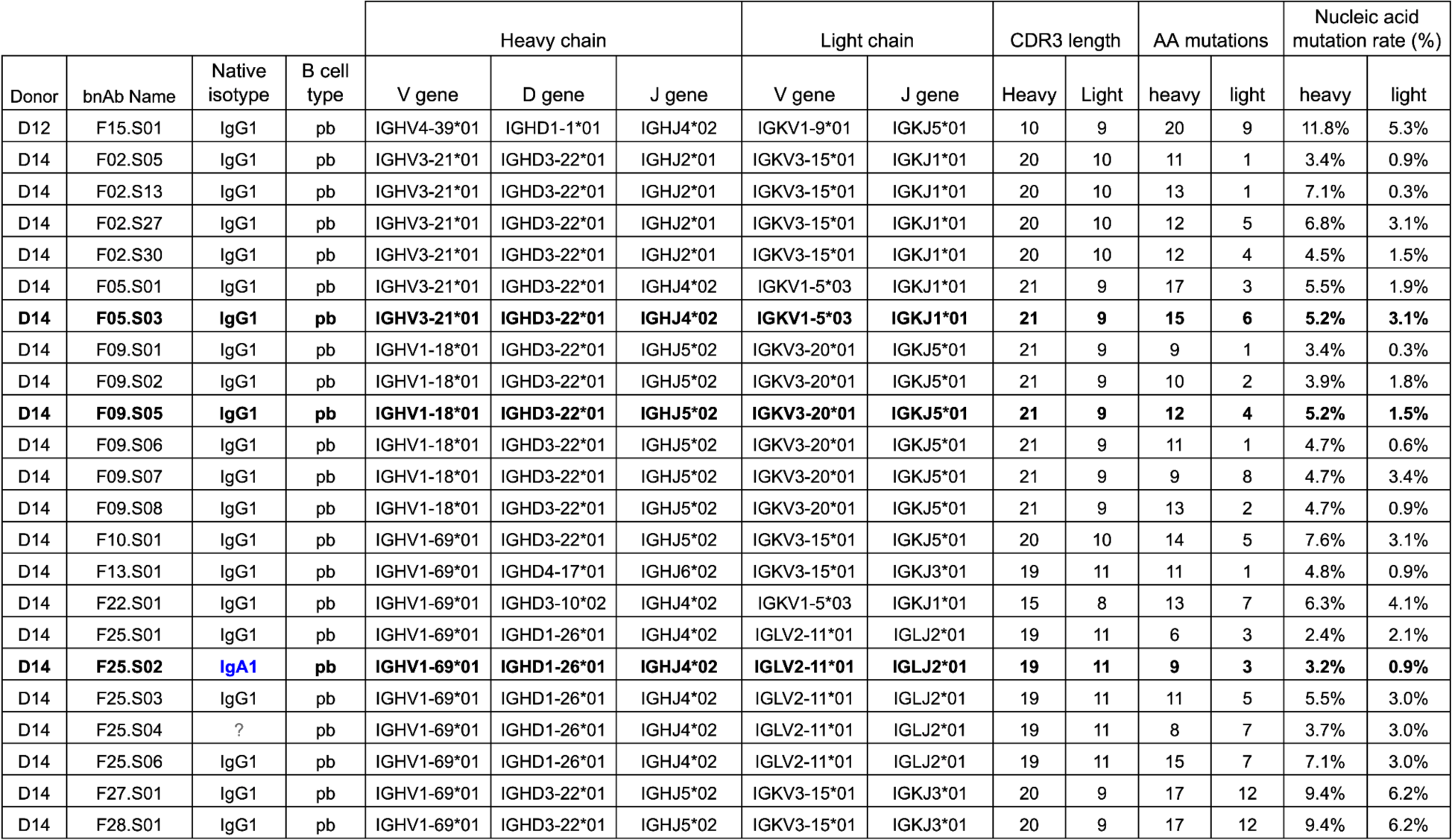
Genetic characteristics of broadly neutralizing antibodies whose IC50 values are displayed in Figure S3. Bold = chosen for detailed characterization; blue = non-IgG isotype;= insufficient sequence coverage of constant gene to determine isotype information; pb = plasmablast.

**Table S3.**
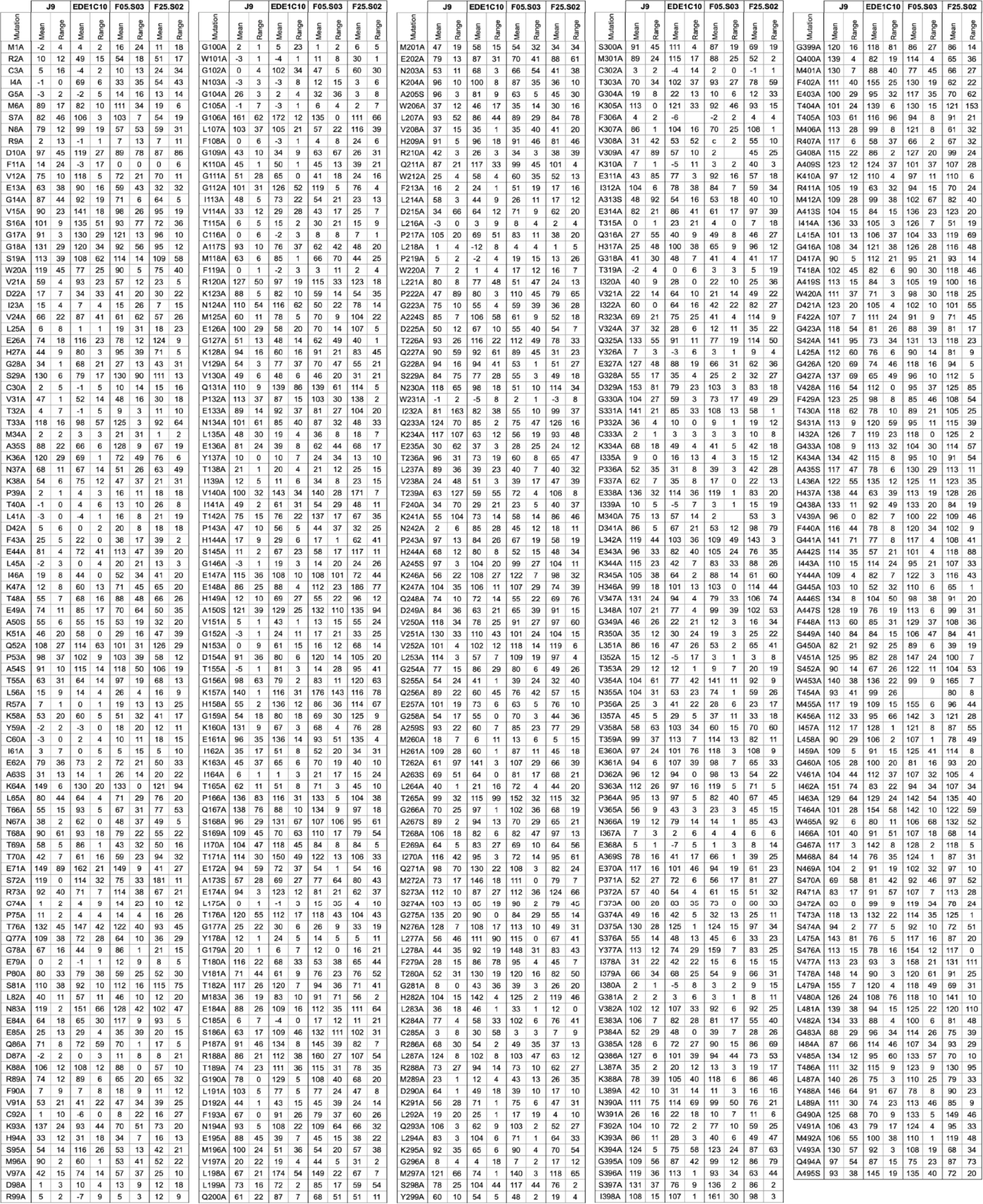
Antibody binding reactivity to DENV2 16681 E protein alanine scanning mutagenesis library. Mean percentage and range of binding reactivity to alanine mutant relative to wild type DENV2 from at least two independent experiments.

**Figure S1.**
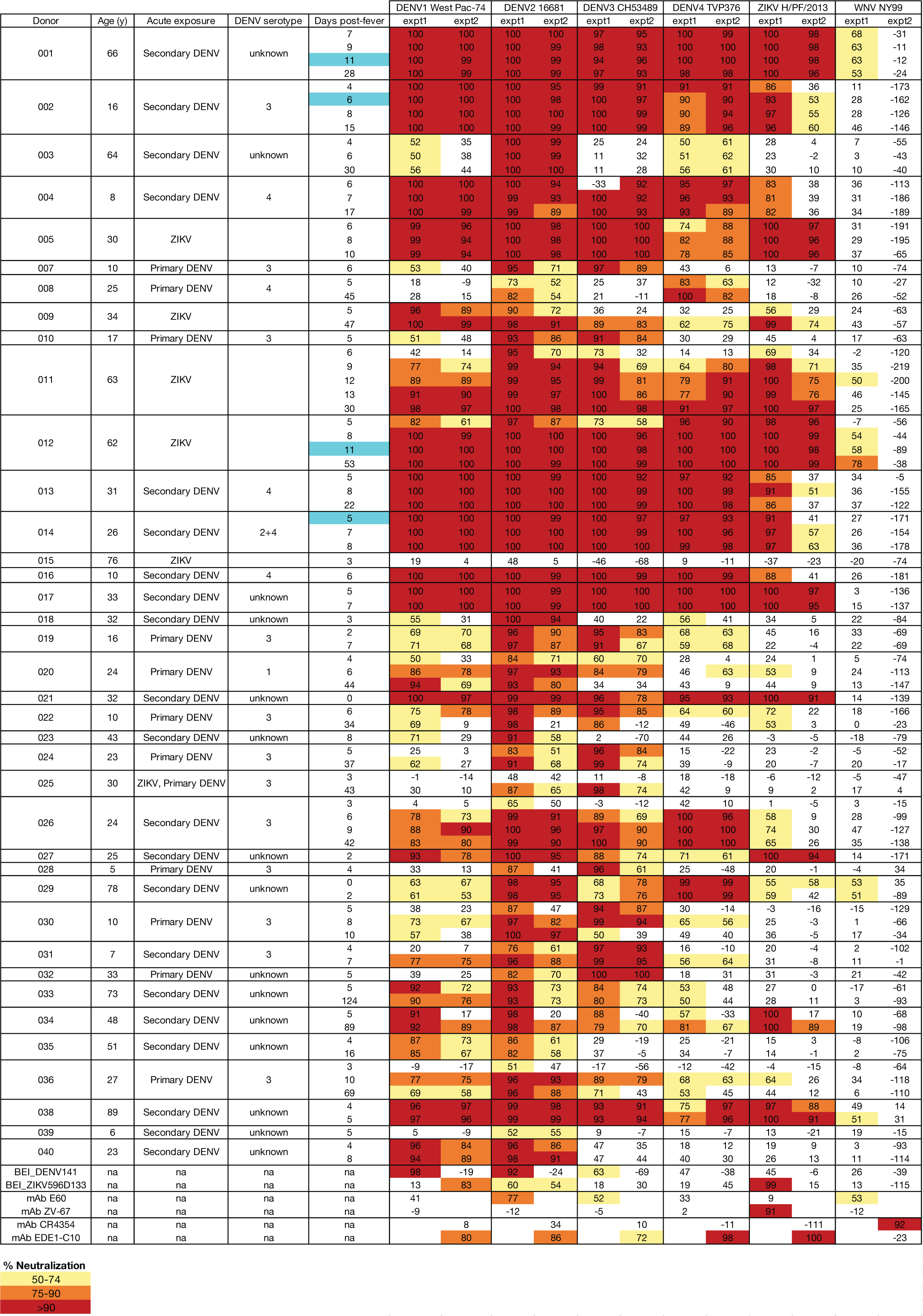
Serum neutralizing activity against flaviviruses. Serum samples from 38 cohort participants with the indicated age and DENV and/or ZIKV exposure histories collected at the time point(s) shown were diluted either 1:240 (expt1) or 1:300 (expt2) and tested for their ability to neutralize the indicated flaviviruses in two independent experiments. Bottom rows indicate control antibodies, which include human convalescent sera to DENV (BEI Resources NR-50232) or ZIKV (BEI Resources NR-50752) and monoclonal antibodies (mAb) E60 ^134^, ZV-67 ^135^, CR4354 ^136^, and EDE1 C10 ^28^. The percent neutralizing activity shown under each virus column is normalized to infection in the absence of antibody. Heatmap colors represent neutralizing activity of at least 50% as indicated in the key under the table. We selected corresponding PBMC samples from the donors and time points highlighted in blue under the ‘Days post-fever’ column for single-cell RNA sequencing to isolate monoclonal antibodies.

**Figure S2.**
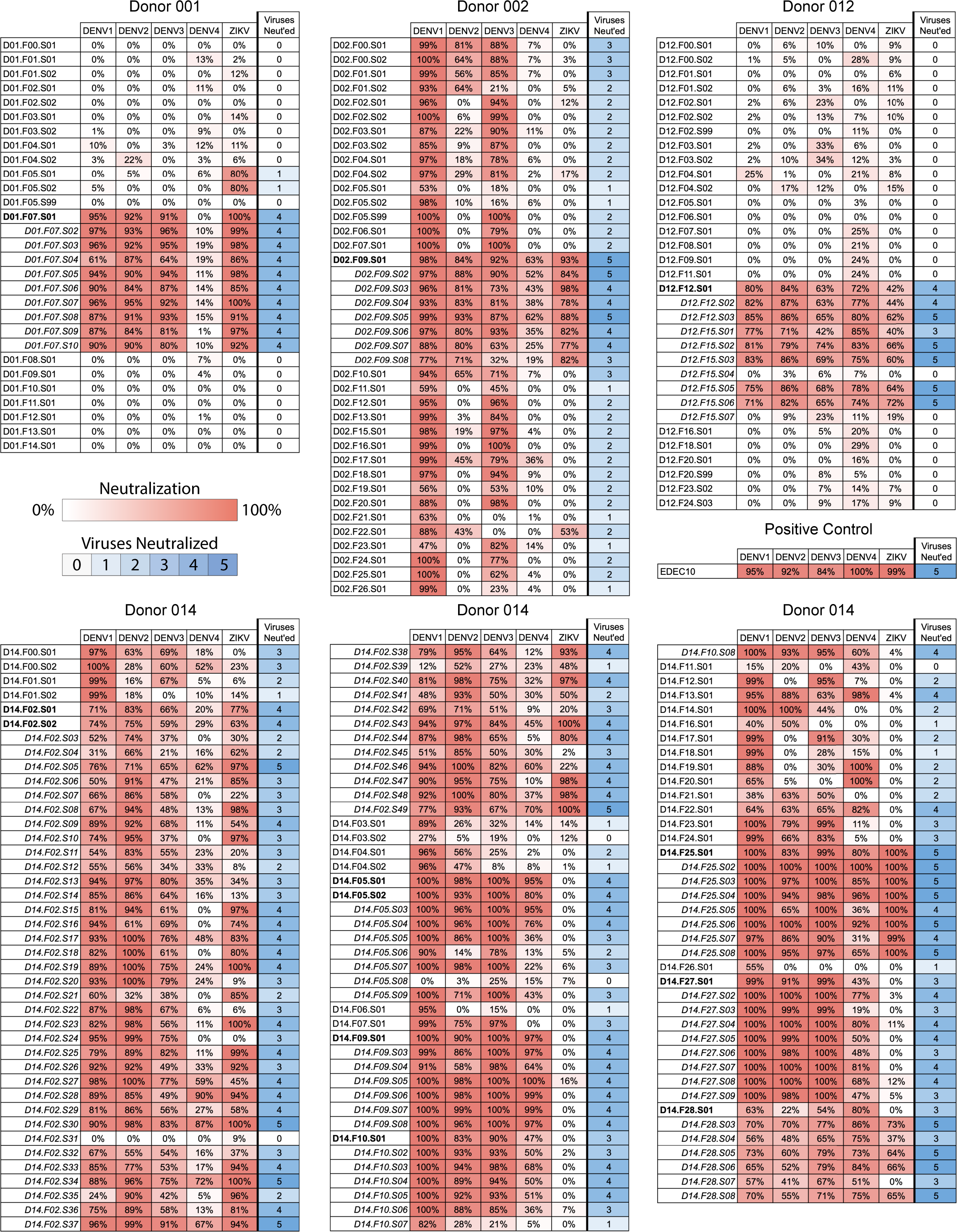
Neutralization profiles of IgG1 transfection supernatants. Heatmaps displaying the results of neutralization assays against DENV1-4 and ZIKV using 1/10 diluted ExpiCHO-S culture supernatant containing the antibodies indicated. Antibodies were named based on the source of the antibody in the format DXX.FYY.SZZ, where XX is the donor number, YY is the clonal family within the donor ranked by decreasing size, and ZZ is assigned by the chronological order in which antibodies from the family were produced. The percent neutralization is calculated relative to infection in the absence of antibody. The final column displays the number of viruses that were neutralized by >50% by that antibody. The antibodies whose names are left aligned were screened in round 1, which was intended to screen many different families. Antibodies that were considered hits due to the breadth and/or potency of their neutralization in round 1 are shown in bold font. For round 2 we selected additional antibodies, shown indented and italicized, from the clonal families of hits identified in round 1. EDE1-C10, which served as a positive control, was expressed and assayed in parallel in every trial of both rounds.

**Figure S3.**
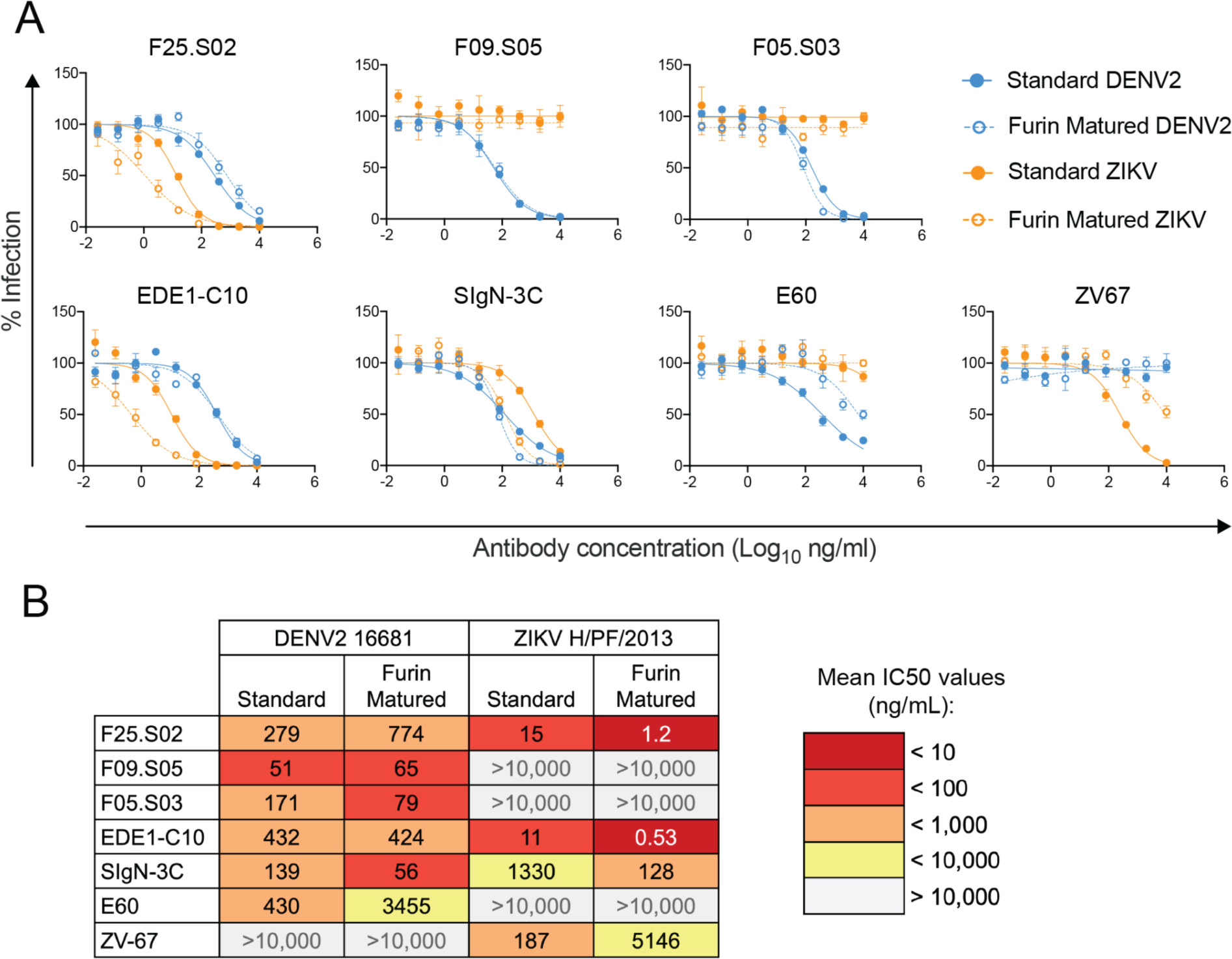
Effect of virion maturation state on bnAb activity. **(A)** The indicated antibodies were tested against DENV2 16681 (blue) or ZIKV H/PF/2013 (orange) reporter virus particles prepared either under standard conditions (solid circles and lines) or in the presence of excess furin (open circles and dashed lines). Data were obtained from two independent experiments, each performed in duplicate wells. Data points and error bars represent the mean infection and standard deviation of the four total replicates, respectively. **(B)** The table displays the mean IC50 values at which the indicated antibodies neutralized the indicated forms of DENV and ZIKV in dose response neutralization curves as shown in (A).

**Figure S4.**
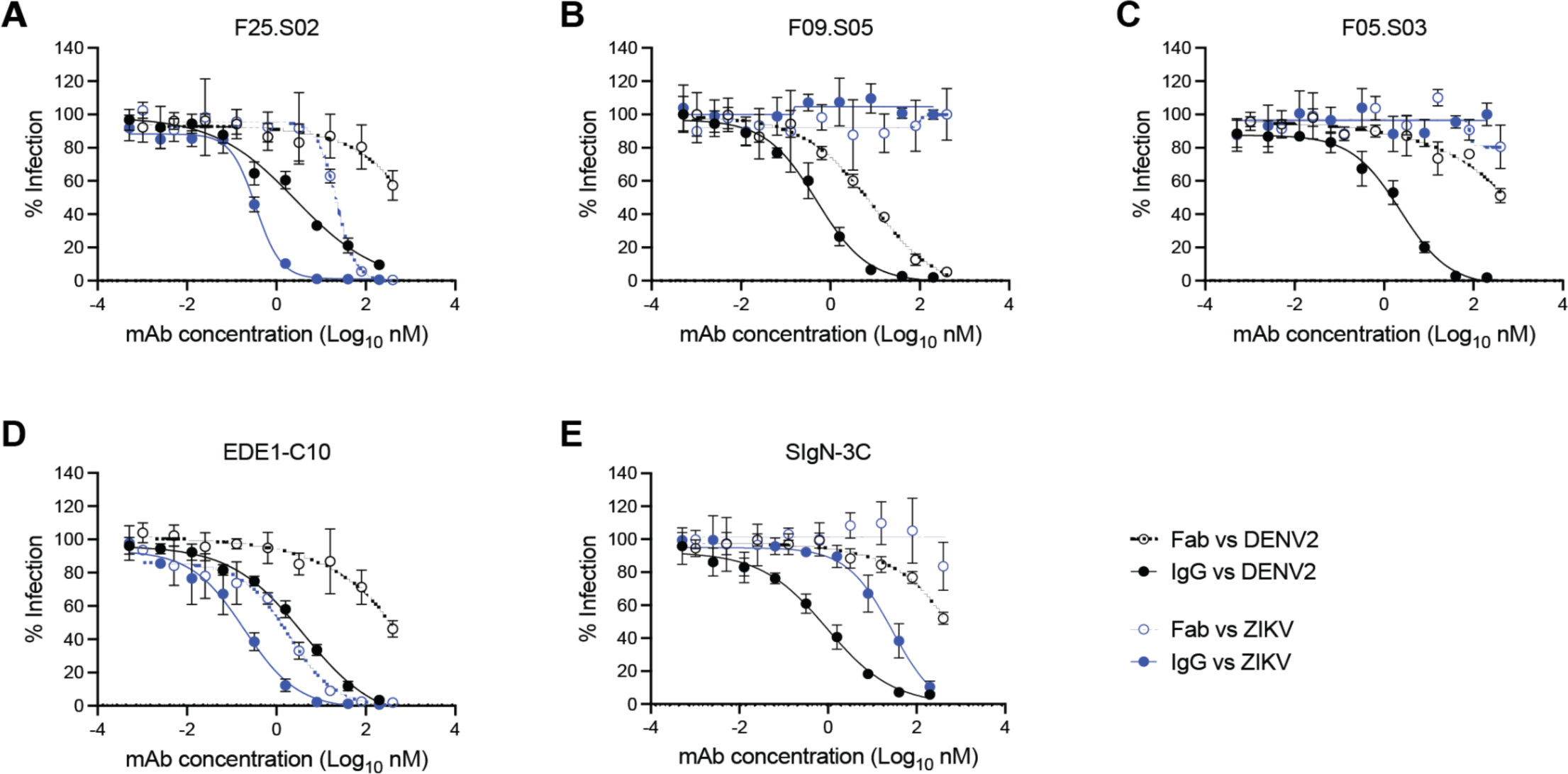
Effect of antibody valency on neutralizing activity. We tested monovalent Fab (open circles and dashed lines) or bivalent IgG (solid circles and lines) versions of antibodies **(A)** F25.S02, **(B)** F09.S05, **(C)** F05.S03, **(D)** EDE1-C10, and **(E)** SIgN-3C against DENV2 16681 (black) or ZIKV H/PF/2013 (blue) reporter virus particles. Dose-response neutralization curves shown are from two independent experiments, each performed in duplicate wells. Data points and error bars represent the mean infection and standard deviation of the four total replicates, respectively.

**Figure S5.**
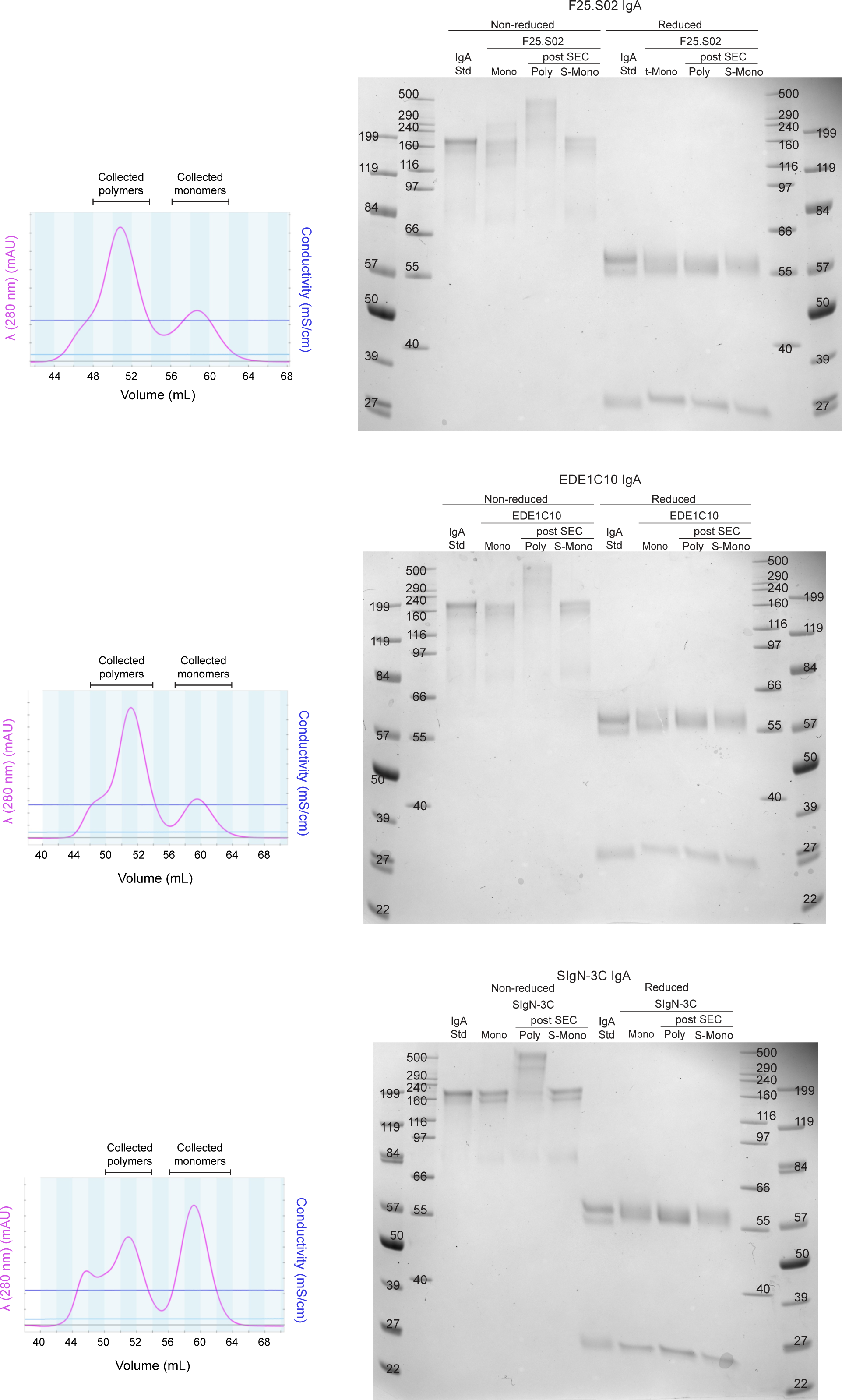
Purity of IgA1 antibody preparations. Graphs on the left display absorbance profiles (at 280 nm) of eluates from size-exclusion chromatography (SEC), which was used to separate monomeric and polymeric IgA1. Images on the right display SDS-PAGE gels to assess purity of preparations. Eluates from SEC were collected in 2 mL fractions and the fractions indicated were collected, pooled, and concentrated to obtain purified monomers and polymers. SDS-PAGE was run on non-reduced (left half) and reduced (right half) samples of each type of antibody. Each half of a gel has one well containing a commercially purchased IgA1 isotype control (IgA Std). Each half also has wells containing two types of IgA1 monomers. The first was produced as monomers, i.e in the absence of a J chain expression plasmid (Mono). The second were produced in a transfection that included a J chain expression plasmid and they were separated from polymers via SEC (S-Mono).

**Figure S6.**
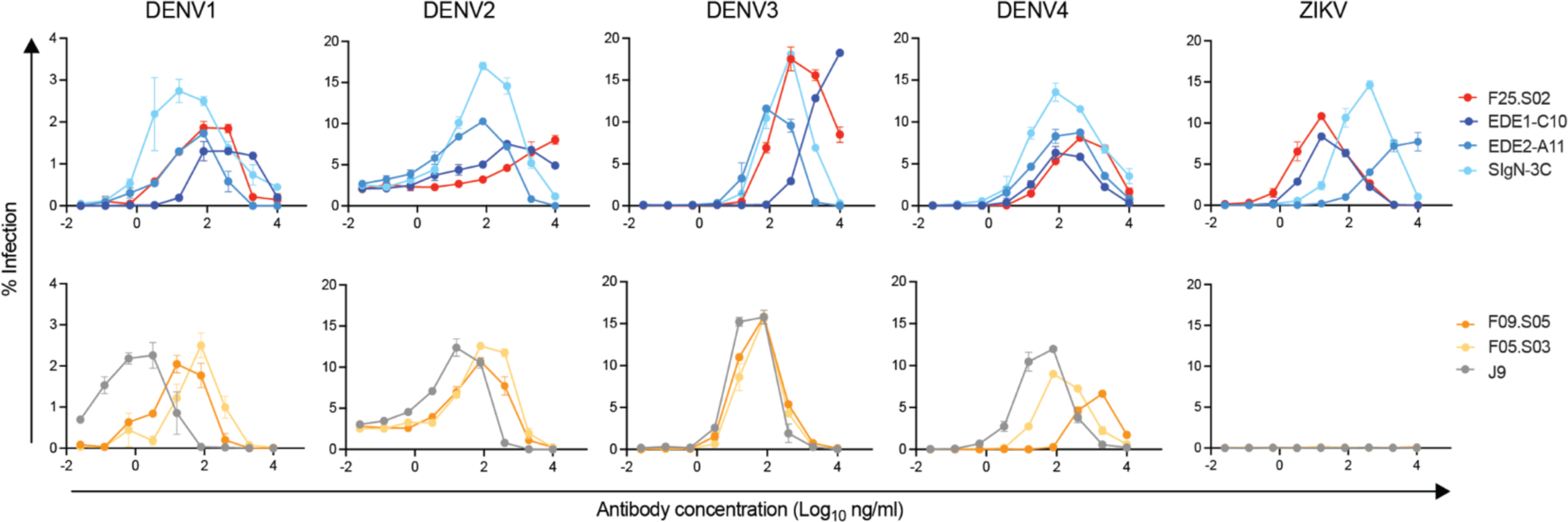
ADE profile of IgG1 bnAbs in K562 cells. Serial dilutions of antibodies indicated in the key were complexed with reporter virus particles shown above graphs prior to infection of K562 cells. Dose-response ADE profiles of antibodies that do or do not neutralize ZIKV in addition to DENV1-4 are shown in top and bottom panels, respectively. Data points and error bars indicate the mean and range of infection in duplicate wells, respectively. Graphs shown are representative of 4-5 independent experiments.

**Figure S7.**
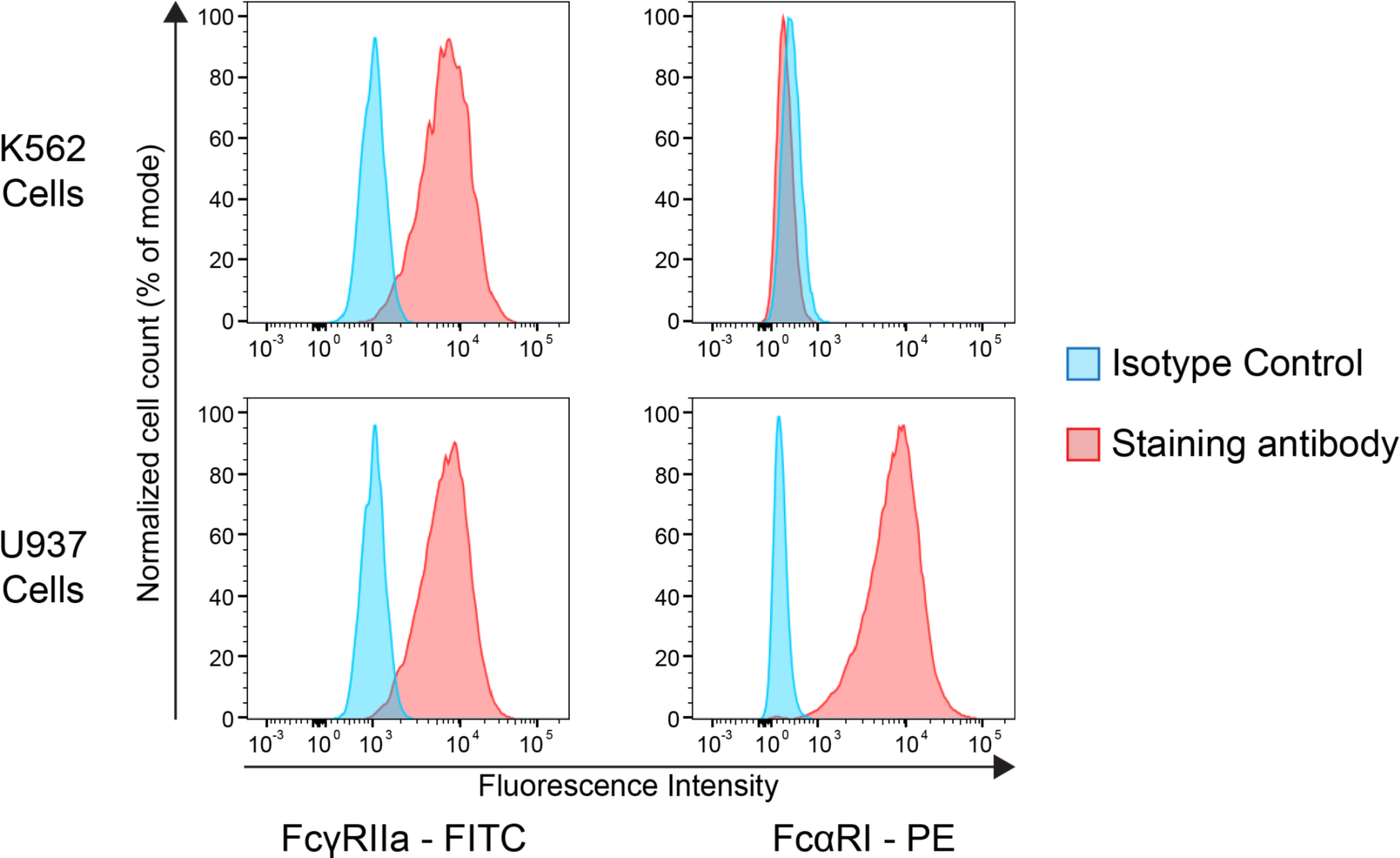
Fc receptor expression profile of K562 and U937 cells. Histograms display the fluorescence intensity of K562 (top row) or U937 (bottom row) cells stained for the indicated Fc receptors. Histograms are normalized to the modal cell count. The isotype control was conjugated to the same fluorophore and used at the same concentration as anti-FcγRIIa or anti-FcαR1 antibody on the same population of cells.

## REFERENCES

1. Pierson, T.C., and Diamond, M.S. (2020). The continued threat of emerging flaviviruses. Nat Microbiol 5, 796–812.

2. Kraemer, M.U.G., Reiner, R.C., Jr, Brady, O.J., Messina, J.P., Gilbert, M., Pigott, D.M., Yi, D., Johnson, K., Earl, L., Marczak, L.B., et al. (2019). Past and future spread of the arbovirus vectors Aedes aegypti and Aedes albopictus. Nat Microbiol 4, 854–863.

3. Messina, J.P., Brady, O.J., Golding, N., Kraemer, M.U.G., Wint, G.R.W., Ray, S.E., Pigott, D.M., Shearer, F.M., Johnson, K., Earl, L., et al. (2019). The current and future global distribution and population at risk of dengue. Nat Microbiol 4, 1508–1515.

4. Iwamura, T., Guzman-Holst, A., and Murray, K.A. (2020). Accelerating invasion potential of disease vector Aedes aegypti under climate change. Nat. Commun. 11, 2130.

5. Waggoner, J.J., Balmaseda, A., Gresh, L., Sahoo, M.K., Montoya, M., Wang, C., Abeynayake, J., Kuan, G., Pinsky, B.A., and Harris, E. (2016). Homotypic Dengue Virus Reinfections in Nicaraguan Children. J. Infect. Dis. 214, 986–993.

6. Forshey, B.M., Reiner, R.C., Olkowski, S., Morrison, A.C., Espinoza, A., Long, K.C., Vilcarromero, S., Casanova, W., Wearing, H.J., Halsey, E.S., et al. (2016). Incomplete Protection against Dengue Virus Type 2 Re-infection in Peru. PLoS Negl. Trop. Dis. 10, e0004398.

7. Snow, G.E., Haaland, B., Ooi, E.E., and Gubler, D.J. (2014). Review article: Research on dengue during World War II revisited. Am. J. Trop. Med. Hyg. 91, 1203–1217.

8. Beltramello, M., Williams, K.L., Simmons, C.P., Macagno, A., Simonelli, L., Quyen, N.T.H., Sukupolvi-Petty, S., Navarro-Sanchez, E., Young, P.R., de Silva, A.M., et al. (2010). The human immune response to Dengue virus is dominated by highly cross-reactive antibodies endowed with neutralizing and enhancing activity. Cell Host Microbe 8, 271–283.

9. de Alwis, R., Beltramello, M., Messer, W.B., Sukupolvi-Petty, S., Wahala, W.M.P.B., Kraus, A., Olivarez, N.P., Pham, Q., Brien, J.D., Tsai, W.-Y., et al. (2011). In-depth analysis of the antibody response of individuals exposed to primary dengue virus infection. PLoS Negl. Trop. Dis. 5, e1188.

10. Lai, C.-Y., Tsai, W.-Y., Lin, S.-R., Kao, C.-L., Hu, H.-P., King, C.-C., Wu, H.-C., Chang, G.-J., and Wang, W.-K. (2008). Antibodies to envelope glycoprotein of dengue virus during the natural course of infection are predominantly cross-reactive and recognize epitopes containing highly conserved residues at the fusion loop of domain II. J. Virol. 82, 6631–6643.

11. Smith, S.A., de Alwis, R., Kose, N., Durbin, A.P., Whitehead, S.S., de Silva, A.M., and Crowe, J.E. (2013). Human Monoclonal Antibodies Derived From Memory B Cells Following Live Attenuated Dengue Virus Vaccination or Natural Infection Exhibit Similar Characteristics. The Journal of Infectious Diseases 207, 1898–1908. 10.1093/infdis/jit119.

12. Katzelnick, L.C., Gresh, L., Halloran, M.E., Mercado, J.C., Kuan, G., Gordon, A., Balmaseda, A., and Harris, E. (2017). Antibody-dependent enhancement of severe dengue disease in humans. Science 358, 929–932.

13. Salje, H., Cummings, D.A.T., Rodriguez-Barraquer, I., Katzelnick, L.C., Lessler, J., Klungthong, C., Thaisomboonsuk, B., Nisalak, A., Weg, A., Ellison, D., et al. (2018). Reconstruction of antibody dynamics and infection histories to evaluate dengue risk. Nature 557, 719–723.

14. Sangkawibha, N., Rojanasuphot, S., Ahandrik, S., Viriyapongse, S., Jatanasen, S., Salitul, V., Phanthumachinda, B., and Halstead, S.B. (1984). Risk factors in dengue shock syndrome: a prospective epidemiologic study in Rayong, Thailand. I. The 1980 outbreak. Am. J. Epidemiol. 120, 653–669.

15. Guzmán, M.G., Kouri, G., Valdes, L., Bravo, J., Alvarez, M., Vazques, S., Delgado, I., and Halstead, S.B. (2000). Epidemiologic studies on Dengue in Santiago de Cuba, 1997. Am. J. Epidemiol. 152, 793–799; discussion 804.

16. Chau, T.N.B., Quyen, N.T.H., Thuy, T.T., Tuan, N.M., Hoang, D.M., Dung, N.T.P., Lien, L.B., Quy, N.T., Hieu, N.T., Hieu, L.T.M., et al. (2008). Dengue in Vietnamese infants--results of infection-enhancement assays correlate with age-related disease epidemiology, and cellular immune responses correlate with disease severity. J. Infect. Dis. 198, 516–524.

17. Wang, T.T., Sewatanon, J., Memoli, M.J., Wrammert, J., Bournazos, S., Bhaumik, S.K., Pinsky, B.A., Chokephaibulkit, K., Onlamoon, N., Pattanapanyasat, K., et al. (2017). IgG antibodies to dengue enhanced for FcγRIIIA binding determine disease severity. Science 355, 395–398.

18. Guzman, M.G., and Harris, E. (2015). Dengue. Lancet 385, 453–465.

19. Hadinegoro, S.R., Arredondo-García, J.L., Capeding, M.R., Deseda, C., Chotpitayasunondh, T., Dietze, R., Muhammad Ismail, H.I.H., Reynales, H., Limkittikul, K., Rivera-Medina, D.M., et al. (2015). Efficacy and Long-Term Safety of a Dengue Vaccine in Regions of Endemic Disease. N. Engl. J. Med. 373, 1195–1206.

20. Villar, L., Dayan, G.H., Arredondo-García, J.L., Rivera, D.M., Cunha, R., Deseda, C., Reynales, H., Costa, M.S., Morales-Ramírez, J.O., Carrasquilla, G., et al. (2015). Efficacy of a tetravalent dengue vaccine in children in Latin America. N. Engl. J. Med. 372, 113–123.

21. Katzelnick, L.C., Narvaez, C., Arguello, S., Lopez Mercado, B., Collado, D., Ampie, O., Elizondo, D., Miranda, T., Bustos Carillo, F., Mercado, J.C., et al. (2020). Zika virus infection enhances future risk of severe dengue disease. Science 369, 1123–1128.

22. Tsai, W.-Y., Lai, C.-Y., Wu, Y.-C., Lin, H.-E., Edwards, C., Jumnainsong, A., Kliks, S., Halstead, S., Mongkolsapaya, J., Screaton, G.R., et al. (2013). High-avidity and potently neutralizing cross-reactive human monoclonal antibodies derived from secondary dengue virus infection. J. Virol. 87, 12562–12575.

23. Lai, C.-Y., Williams, K.L., Wu, Y.-C., Knight, S., Balmaseda, A., Harris, E., and Wang, W.-K. (2013). Analysis of cross-reactive antibodies recognizing the fusion loop of envelope protein and correlation with neutralizing antibody titers in Nicaraguan dengue cases. PLoS Negl. Trop. Dis. 7, e2451.

24. Andrade, P., Narvekar, P., Montoya, M., Michlmayr, D., Balmaseda, A., Coloma, J., and Harris, E. (2020). Primary and Secondary Dengue Virus Infections Elicit Similar Memory B-Cell Responses, but Breadth to Other Serotypes and Cross-Reactivity to Zika Virus Is Higher in Secondary Dengue. J. Infect. Dis. 222, 590–600.

25. Zompi, S., Montoya, M., Pohl, M.O., Balmaseda, A., and Harris, E. (2012). Dominant cross-reactive B cell response during secondary acute dengue virus infection in humans. PLoS Negl. Trop. Dis. 6, e1568.

26. Gibbons, R.V., Kalanarooj, S., Jarman, R.G., Nisalak, A., Vaughn, D.W., Endy, T.P., Mammen, M.P., Jr, and Srikiatkhachorn, A. (2007). Analysis of repeat hospital admissions for dengue to estimate the frequency of third or fourth dengue infections resulting in admissions and dengue hemorrhagic fever, and serotype sequences. Am. J. Trop. Med. Hyg. 77, 910–913.

27. Xu, M., Zuest, R., Velumani, S., Tukijan, F., Toh, Y.X., Appanna, R., Tan, E.Y., Cerny, D., MacAry, P., Wang, C.-I., et al. (2017). A potent neutralizing antibody with therapeutic potential against all four serotypes of dengue virus. NPJ Vaccines 2, 2.

28. Dejnirattisai, W., Wongwiwat, W., Supasa, S., Zhang, X., Dai, X., Rouvinski, A., Jumnainsong, A., Edwards, C., Quyen, N.T.H., Duangchinda, T., et al. (2015). A new class of highly potent, broadly neutralizing antibodies isolated from viremic patients infected with dengue virus. Nat. Immunol. 16, 170–177.

29. Smith, S.A., de Alwis, A.R., Kose, N., Harris, E., Ibarra, K.D., Kahle, K.M., Pfaff, J.M., Xiang, X., Doranz, B.J., de Silva, A.M., et al. (2013). The potent and broadly neutralizing human dengue virus-specific monoclonal antibody 1C19 reveals a unique cross-reactive epitope on the bc loop of domain II of the envelope protein. MBio 4, e00873–13.

30. Rouvinski, A., Guardado-Calvo, P., Barba-Spaeth, G., Duquerroy, S., Vaney, M.-C., Kikuti, C.M., Navarro Sanchez, M.E., Dejnirattisai, W., Wongwiwat, W., Haouz, A., et al. (2015). Recognition determinants of broadly neutralizing human antibodies against dengue viruses. Nature 520, 109– 113.

31. Barba-Spaeth, G., Dejnirattisai, W., Rouvinski, A., Vaney, M.-C., Medits, I., Sharma, A., Simon-Lorière, E., Sakuntabhai, A., Cao-Lormeau, V.-M., Haouz, A., et al. (2016). Structural basis of potent Zika-dengue virus antibody cross-neutralization. Nature 536, 48–53.

32. Kotaki, T., Kurosu, T., Grinyo-Escuer, A., Davidson, E., Churrotin, S., Okabayashi, T., Puiprom, O., Mulyatno, K.C., Sucipto, T.H., Doranz, B.J., et al. (2021). An affinity-matured human monoclonal antibody targeting fusion loop epitope of dengue virus with in vivo therapeutic potency. Sci. Rep. 11, 12987.

33. Dussupt, V., Sankhala, R.S., Gromowski, G.D., Donofrio, G., De La Barrera, R.A., Larocca, R.A., Zaky, W., Mendez-Rivera, L., Choe, M., Davidson, E., et al. (2020). Potent Zika and dengue cross-neutralizing antibodies induced by Zika vaccination in a dengue-experienced donor. Nat. Med. 26, 228–235.

34. Robbiani, D.F., Bozzacco, L., Keeffe, J.R., Khouri, R., Olsen, P.C., Gazumyan, A., Schaefer-Babajew, D., Avila-Rios, S., Nogueira, L., Patel, R., et al. (2017). Recurrent Potent Human Neutralizing Antibodies to Zika Virus in Brazil and Mexico. Cell 169, 597–609.e11.

35. Rogers, T.F., Goodwin, E.C., Briney, B., Sok, D., Beutler, N., Strubel, A., Nedellec, R., Le, K., Brown, M.E., Burton, D.R., et al. (2017). Zika virus activates de novo and cross-reactive memory B cell responses in dengue-experienced donors. Sci Immunol 2. 10.1126/sciimmunol.aan6809.

36. Kam, Y.-W., Lee, C.Y.-P., Teo, T.-H., Howland, S.W., Amrun, S.N., Lum, F.-M., See, P., Kng, N.Q.-R., Huber, R.G., Xu, M.-H., et al. (2017). Cross-reactive dengue human monoclonal antibody prevents severe pathologies and death from Zika virus infections. JCI Insight 2. 10.1172/jci.insight.92428.

37. Zhang, S., Loy, T., Ng, T.-S., Lim, X.-N., Chew, S.-Y.V., Tan, T.Y., Xu, M., Kostyuchenko, V.A., Tukijan, F., Shi, J., et al. (2020). A Human Antibody Neutralizes Different Flaviviruses by Using Different Mechanisms. Cell Rep. 31, 107584.

38. Boonyaratanakornkit, J., and Taylor, J.J. (2019). Techniques to Study Antigen-Specific B Cell Responses. Front. Immunol. 10, 1694.

39. Durham, N.D., Agrawal, A., Waltari, E., Croote, D., Zanini, F., Fouch, M., Davidson, E., Smith, O., Carabajal, E., Pak, J.E., et al. (2019). Broadly neutralizing human antibodies against dengue virus identified by single B cell transcriptomics. Elife 8. 10.7554/eLife.52384.

40. Xu, M., Hadinoto, V., Appanna, R., Joensson, K., Toh, Y.X., Balakrishnan, T., Ong, S.H., Warter, L., Leo, Y.S., Wang, C.-I., et al. (2012). Plasmablasts generated during repeated dengue infection are virus glycoprotein-specific and bind to multiple virus serotypes. J. Immunol. 189, 5877–5885.

41. Zanini, F., Robinson, M.L., Croote, D., Sahoo, M.K., Sanz, A.M., Ortiz-Lasso, E., Albornoz, L.L., Rosso, F., Montoya, J.G., Goo, L., et al. (2018). Virus-inclusive single-cell RNA sequencing reveals the molecular signature of progression to severe dengue. Proc. Natl. Acad. Sci. U. S. A. 115, E12363–E12369.

42. Robinson, M., Sweeney, T.E., Barouch-Bentov, R., Sahoo, M.K., Kalesinskas, L., Vallania, F., Sanz, A.M., Ortiz-Lasso, E., Albornoz, L.L., Rosso, F., et al. (2019). A 20-Gene Set Predictive of Progression to Severe Dengue. Cell Rep. 26, 1104–1111.e4.

43. Zhang, B., Pinsky, B.A., Ananta, J.S., Zhao, S., Arulkumar, S., Wan, H., Sahoo, M.K., Abeynayake, J., Waggoner, J.J., Hopes, C., et al. (2017). Diagnosis of Zika virus infection on a nanotechnology platform. Nat. Med. 23, 548–550.

44. Priyamvada, L., Cho, A., Onlamoon, N., Zheng, N.-Y., Huang, M., Kovalenkov, Y., Chokephaibulkit, K., Angkasekwinai, N., Pattanapanyasat, K., Ahmed, R., et al. (2016). B Cell Responses during Secondary Dengue Virus Infection Are Dominated by Highly Cross-Reactive, Memory-Derived Plasmablasts. J. Virol. 90, 5574–5585.

45. Wrammert, J., Onlamoon, N., Akondy, R.S., Perng, G.C., Polsrila, K., Chandele, A., Kwissa, M., Pulendran, B., Wilson, P.C., Wittawatmongkol, O., et al. (2012). Rapid and massive virus-specific plasmablast responses during acute dengue virus infection in humans. J. Virol. 86, 2911–2918.

46. Nivarthi, U.K., Tu, H.A., Delacruz, M.J., Swanstrom, J., Patel, B., Durbin, A.P., Whitehead, S.S., Pierce, K.K., Kirkpatrick, B.D., Baric, R.S., et al. (2019). Longitudinal analysis of acute and convalescent B cell responses in a human primary dengue serotype 2 infection model. EBioMedicine 41, 465–478.

47. Garcia-Bates, T.M., Cordeiro, M.T., Nascimento, E.J.M., Smith, A.P., Soares de Melo, K.M., McBurney, S.P., Evans, J.D., Marques, E.T.A., Jr, and Barratt-Boyes, S.M. (2013). Association between magnitude of the virus-specific plasmablast response and disease severity in dengue patients. J. Immunol. 190, 80–87.

48. Waickman, A.T., Gromowski, G.D., Rutvisuttinunt, W., Li, T., Siegfried, H., Victor, K., Kuklis, C., Gomootsukavadee, M., McCracken, M.K., Gabriel, B., et al. (2020). Transcriptional and clonal characterization of B cell plasmablast diversity following primary and secondary natural DENV infection. EBioMedicine 54, 102733.

49. Setliff, I., Shiakolas, A.R., Pilewski, K.A., Murji, A.A., Mapengo, R.E., Janowska, K., Richardson, S., Oosthuysen, C., Raju, N., Ronsard, L., et al. (2019). High-Throughput Mapping of B Cell Receptor Sequences to Antigen Specificity. Cell 179, 1636–1646.e15.

50. Cao, Y., Su, B., Guo, X., Sun, W., Deng, Y., Bao, L., Zhu, Q., Zhang, X., Zheng, Y., Geng, C., et al. (2020). Potent Neutralizing Antibodies against SARS-CoV-2 Identified by High-Throughput Single-Cell Sequencing of Convalescent Patients’ B Cells. Cell 182, 73–84.e16.

51. Ralph, D.K., and Matsen, F.A., 4th (2022). Inference of B cell clonal families using heavy/light chain pairing information. PLoS Comput. Biol. 18, e1010723.

52. Croote, D., Darmanis, S., Nadeau, K.C., and Quake, S.R. (2018). High-affinity allergen-specific human antibodies cloned from single IgE B cell transcriptomes. Science 362, 1306–1309.

53. Ralph, D.K., and Matsen, F.A., 4th (2020). Using B cell receptor lineage structures to predict affinity. PLoS Comput. Biol. 16, e1008391.

54. Wang, Q., Michailidis, E., Yu, Y., Wang, Z., Hurley, A.M., Oren, D.A., Mayer, C.T., Gazumyan, A., Liu, Z., Zhou, Y., et al. (2020). A Combination of Human Broadly Neutralizing Antibodies against Hepatitis B Virus HBsAg with Distinct Epitopes Suppresses Escape Mutations. Cell Host Microbe 28, 335–349.e6.

55. Kwong, P.D., and Mascola, J.R. (2012). Human antibodies that neutralize HIV-1: identification, structures, and B cell ontogenies. Immunity 37, 412–425.

56. Katzelnick, L.C., Fonville, J.M., Gromowski, G.D., Bustos Arriaga, J., Green, A., James, S.L., Lau, L., Montoya, M., Wang, C., VanBlargan, L.A., et al. (2015). Dengue viruses cluster antigenically but not as discrete serotypes. Science 349, 1338–1343.

57. Bell, S.M., Katzelnick, L., and Bedford, T. (2019). Dengue genetic divergence generates within-serotype antigenic variation, but serotypes dominate evolutionary dynamics. Elife 8. 10.7554/eLife.42496.

58. Martinez, D.R., Yount, B., Nivarthi, U., Munt, J.E., Delacruz, M.J., Whitehead, S.S., Durbin, A.P., de Silva, A.M., and Baric, R.S. (2020). Antigenic Variation of the Dengue Virus 2 Genotypes Impacts the Neutralization Activity of Human Antibodies in Vaccinees. Cell Rep. 33, 108226.

59. Rico-Hesse, R. (1990). Molecular evolution and distribution of dengue viruses type 1 and 2 in nature. Virology 174, 479–493.

60. Holmes, E.C., and Twiddy, S.S. (2003). The origin, emergence and evolutionary genetics of dengue virus. Infect. Genet. Evol. 3, 19–28.

61. Dowd, K.A., DeMaso, C.R., and Pierson, T.C. (2015). Genotypic Differences in Dengue Virus Neutralization Are Explained by a Single Amino Acid Mutation That Modulates Virus Breathing. MBio 6, e01559–15.

62. VanBlargan, L.A., Milutinovic, P.S., Goo, L., DeMaso, C.R., Durbin, A.P., Whitehead, S.S., Pierson, T.C., and Dowd, K.A. (2021). Dengue Virus Serotype 1 Conformational Dynamics Confers Virus Strain-Dependent Patterns of Neutralization by Polyclonal Sera. J. Virol. 95, e0095621.

63. Chen, R., and Vasilakis, N. (2011). Dengue--quo tu et quo vadis? Viruses 3, 1562–1608.

64. Gallichotte, E.N., Baric, T.J., Nivarthi, U., Delacruz, M.J., Graham, R., Widman, D.G., Yount, B.L., Durbin, A.P., Whitehead, S.S., de Silva, A.M., et al. (2018). Genetic Variation between Dengue Virus Type 4 Strains Impacts Human Antibody Binding and Neutralization. Cell Rep. 25, 1214– 1224.

65. Henein, S., Swanstrom, J., Byers, A.M., Moser, J.M., Shaik, S.F., Bonaparte, M., Jackson, N., Guy, B., Baric, R., and de Silva, A.M. (2017). Dissecting Antibodies Induced by a Chimeric Yellow Fever-Dengue, Live-Attenuated, Tetravalent Dengue Vaccine (CYD-TDV) in Naive and Dengue-Exposed Individuals. J. Infect. Dis. 215, 351–358.

66. Juraska, M., Magaret, C.A., Shao, J., Carpp, L.N., Fiore-Gartland, A.J., Benkeser, D., Girerd-Chambaz, Y., Langevin, E., Frago, C., Guy, B., et al. (2018). Viral genetic diversity and protective efficacy of a tetravalent dengue vaccine in two phase 3 trials. Proc. Natl. Acad. Sci. U. S. A. 115, E8378–E8387.

67. Rabaa, M.A., Girerd-Chambaz, Y., Duong Thi Hue, K., Vu Tuan, T., Wills, B., Bonaparte, M., van der Vliet, D., Langevin, E., Cortes, M., Zambrano, B., et al. (2017). Genetic epidemiology of dengue viruses in phase III trials of the CYD tetravalent dengue vaccine and implications for efficacy. Elife 6. 10.7554/eLife.24196.

68. Normile, D. (2013). Tropical medicine. Surprising new dengue virus throws a spanner in disease control efforts. Science 342, 415.

69. Young, K.I., Mundis, S., Widen, S.G., Wood, T.G., Tesh, R.B., Cardosa, J., Vasilakis, N., Perera, D., and Hanley, K.A. (2017). Abundance and distribution of sylvatic dengue virus vectors in three different land cover types in Sarawak, Malaysian Borneo. Parasit. Vectors 10, 406.

70. Chen, R.E., Smith, B.K., Errico, J.M., Gordon, D.N., Winkler, E.S., VanBlargan, L.A., Desai, C., Handley, S.A., Dowd, K.A., Amaro-Carambot, E., et al. (2021). Implications of a highly divergent dengue virus strain for cross-neutralization, protection, and vaccine immunity. Cell Host Microbe 29, 1634–1648.e5.

71. Cherrier, M.V., Kaufmann, B., Nybakken, G.E., Lok, S.-M., Warren, J.T., Chen, B.R., Nelson, C.A., Kostyuchenko, V.A., Holdaway, H.A., Chipman, P.R., et al. (2009). Structural basis for the preferential recognition of immature flaviviruses by a fusion-loop antibody. EMBO J. 28, 3269– 3276.

72. Nelson, S., Jost, C.A., Xu, Q., Ess, J., Martin, J.E., Oliphant, T., Whitehead, S.S., Durbin, A.P., Graham, B.S., Diamond, M.S., et al. (2008). Maturation of West Nile virus modulates sensitivity to antibody-mediated neutralization. PLoS Pathog. 4, e1000060.

73. Goo, L., Debbink, K., Kose, N., Sapparapu, G., Doyle, M.P., Wessel, A.W., Richner, J.M., Burgomaster, K.E., Larman, B.C., Dowd, K.A., et al. (2019). A protective human monoclonal antibody targeting the West Nile virus E protein preferentially recognizes mature virions. Nat Microbiol 4, 71–77.

74. Raut, R., Corbett, K.S., Tennekoon, R.N., Premawansa, S., Wijewickrama, A., Premawansa, G., Mieczkowski, P., Rückert, C., Ebel, G.D., De Silva, A.D., et al. (2019). Dengue type 1 viruses circulating in humans are highly infectious and poorly neutralized by human antibodies. Proc. Natl. Acad. Sci. U. S. A. 116, 227–232.

75. Maciejewski, S., Ruckwardt, T.J., Morabito, K.M., Foreman, B.M., Burgomaster, K.E., Gordon, D.N., Pelc, R.S., DeMaso, C.R., Ko, S.-Y., Fisher, B.E., et al. (2020). Distinct neutralizing antibody correlates of protection among related Zika virus vaccines identify a role for antibody quality. Sci. Transl. Med. 12. 10.1126/scitranslmed.aaw9066.

76. Goo, L., VanBlargan, L.A., Dowd, K.A., Diamond, M.S., and Pierson, T.C. (2017). A single mutation in the envelope protein modulates flavivirus antigenicity, stability, and pathogenesis. PLoS Pathog. 13, e1006178.

77. Goo, L., DeMaso, C.R., Pelc, R.S., Ledgerwood, J.E., Graham, B.S., Kuhn, R.J., and Pierson, T.C. (2018). The Zika virus envelope protein glycan loop regulates virion antigenicity. Virology 515, 191–202.

78. VanBlargan, L.A., Goo, L., and Pierson, T.C. (2016). Deconstructing the Antiviral Neutralizing-Antibody Response: Implications for Vaccine Development and Immunity. Microbiol. Mol. Biol. Rev. 80, 989–1010.

79. Davidson, E., and Doranz, B.J. (2014). A high-throughput shotgun mutagenesis approach to mapping B-cell antibody epitopes. Immunology 143, 13–20.

80. Sharma, A., Zhang, X., Dejnirattisai, W., Dai, X., Gong, D., Wongwiwat, W., Duquerroy, S., Rouvinski, A., Vaney, M.-C., Guardado-Calvo, P., et al. (2021). The epitope arrangement on flavivirus particles contributes to Mab C10’s extraordinary neutralization breadth across Zika and dengue viruses. Cell 184, 6052–6066.e18.

81. Scheepers, C., Richardson, S.I., Moyo-Gwete, T., and Moore, P.L. (2022). Antibody class-switching as a strategy to improve HIV-1 neutralization. Trends Mol. Med. 28, 979–988.

82. Scheepers, C., Bekker, V., Anthony, C., Richardson, S.I., Oosthuysen, B., Moyo, T., Kgagudi, P., Kitchin, D., Nonyane, M., York, T., et al. (2020). Antibody Isotype Switching as a Mechanism to Counter HIV Neutralization Escape. Cell Rep. 33, 108430.

83. Wang, Z., Lorenzi, J.C.C., Muecksch, F., Finkin, S., Viant, C., Gaebler, C., Cipolla, M., Hoffmann, H.-H., Oliveira, T.Y., Oren, D.A., et al. (2021). Enhanced SARS-CoV-2 neutralization by dimeric IgA. Sci. Transl. Med. 13. 10.1126/scitranslmed.abf1555.

84. Bolton, M.J., Arevalo, C.P., Griesman, T., Li, S.H., Bates, P., Wilson, P.C., and Hensley, S.E. (2022). IgG3 subclass antibodies recognize antigenically drifted influenza viruses and SARS-CoV-2 variants through efficient bivalent binding. bioRxiv. 10.1101/2022.09.27.509738.

85. Hale, M., Netland, J., Chen, Y., Thouvenel, C.D., Smith, K.N., Rich, L.M., Vanderwall, E.R., Miranda, M.C., Eggenberger, J., Hao, L., et al. (2022). IgM antibodies derived from memory B cells are potent cross-variant neutralizers of SARS-CoV-2. J. Exp. Med. 219. 10.1084/jem.20220849.

86. Singh, T., Hwang, K.-K., Miller, A.S., Jones, R.L., Lopez, C.A., Dulson, S.J., Giuberti, C., Gladden, M.A., Miller, I., Webster, H.S., et al. (2022). A Zika virus-specific IgM elicited in pregnancy exhibits ultrapotent neutralization. Cell 185, 4826–4840.e17.

87. Woof, J.M., and Russell, M.W. (2011). Structure and function relationships in IgA. Mucosal Immunol. 4, 590–597.

88. Pierson, T.C., Xu, Q., Nelson, S., Oliphant, T., Nybakken, G.E., Fremont, D.H., and Diamond, M.S. (2007). The stoichiometry of antibody-mediated neutralization and enhancement of West Nile virus infection. Cell Host Microbe 1, 135–145.

89. Littaua, R., Kurane, I., and Ennis, F.A. (1990). Human IgG Fc receptor II mediates antibody-dependent enhancement of dengue virus infection. J. Immunol. 144, 3183–3186.

90. Thulin, N.K., Brewer, R.C., Sherwood, R., Bournazos, S., Edwards, K.G., Ramadoss, N.S., Taubenberger, J.K., Memoli, M., Gentles, A.J., Jagannathan, P., et al. (2020). Maternal Anti-Dengue IgG Fucosylation Predicts Susceptibility to Dengue Disease in Infants. Cell Rep. 31, 107642.

91. Bournazos, S., Gupta, A., and Ravetch, J.V. (2020). The role of IgG Fc receptors in antibody-dependent enhancement. Nat. Rev. Immunol. 20, 633–643.

92. Arvin, A.M., Fink, K., Schmid, M.A., Cathcart, A., Spreafico, R., Havenar-Daughton, C., Lanzavecchia, A., Corti, D., and Virgin, H.W. (2020). A perspective on potential antibody-dependent enhancement of SARS-CoV-2. Nature 584, 353–363.

93. Boltz-Nitulescu, G., Willheim, M., Spittler, A., Leutmezer, F., Tempfer, C., and Winkler, S. (1995). Modulation of IgA, IgE, and IgG Fc receptor expression on human mononuclear phagocytes by 1 alpha,25-dihydroxyvitamin D3 and cytokines. J. Leukoc. Biol. 58, 256–262.

94. Geissmann, F., Launay, P., Pasquier, B., Lepelletier, Y., Leborgne, M., Lehuen, A., Brousse, N., and Monteiro, R.C. (2001). A subset of human dendritic cells expresses IgA Fc receptor (CD89), which mediates internalization and activation upon cross-linking by IgA complexes. J. Immunol. 166, 346–352.

95. Parola, C., Neumeier, D., and Reddy, S.T. (2018). Integrating high-throughput screening and sequencing for monoclonal antibody discovery and engineering. Immunology 153, 31–41.

96. Gilchuk, P., Bombardi, R.G., Erasmus, J.H., Tan, Q., Nargi, R., Soto, C., Abbink, P., Suscovich, T.J., Durnell, L.A., Khandhar, A., et al. (2020). Integrated pipeline for the accelerated discovery of antiviral antibody therapeutics. Nat Biomed Eng 4, 1030–1043.

97. Senaratne, U.T.N., Murugananthan, K., Sirisena, P.D.N.N., Carr, J.M., and Noordeen, F. (2020). Dengue virus co-infections with multiple serotypes do not result in a different clinical outcome compared to mono-infections. Epidemiol. Infect. 148, e119.

98. Gubler, D.J., Kuno, G., Sather, G.E., and Waterman, S.H. (1985). A case of natural concurrent human infection with two dengue viruses. Am. J. Trop. Med. Hyg. 34, 170–173.

99. Bharaj, P., Chahar, H.S., Pandey, A., Diddi, K., Dar, L., Guleria, R., Kabra, S.K., and Broor, S. (2008). Concurrent infections by all four dengue virus serotypes during an outbreak of dengue in 2006 in Delhi, India. Virology Journal 5. 10.1186/1743-422x-5-1.

100. Figueiredo, R.M.P. de, Naveca, F.G., Oliveira, C.M., Bastos, M. de S., Mourão, M.P.G., Viana, S. de S., Melo, M. do N., Itapirema, E.F., Saatkamp, C.J., and Farias, I.P. (2011). Co-infection of Dengue virus by serotypes 3 and 4 in patients from Amazonas, Brazil. Rev. Inst. Med. Trop. Sao Paulo 53, 321–323.

101. Colombo, T.E., Vedovello, D., Mondini, A., Reis, A.F.N., Cury, A.A.F., de Oliveira, F.H., Cruz, L.E.A., de Morais Bronzoni, R.V., and Nogueira, M.L. (2013). CO-INFECTION OF DENGUE VIRUS BY SEROTYPES 1 AND 4 IN PATIENT FROM MEDIUM SIZED CITY FROM BRAZIL. Revista do Instituto de Medicina Tropical de São Paulo 55, 275–281. 10.1590/s0036-46652013000400009.

102. Dewi, B.E., Naiggolan, L., Putri, D.H., Rachmayanti, N., Albar, S., Indriastuti, N.T., Sjamsuridzal, W., and Sudiro, T.M. (2014). Characterization of dengue virus serotype 4 infection in Jakarta, Indonesia. Southeast Asian J. Trop. Med. Public Health 45, 53–61.

103. Neumeier, D., Yermanos, A., Agrafiotis, A., Csepregi, L., Chowdhury, T., Ehling, R.A., Kuhn, R., Cotet, T.-S., Brisset-Di Roberto, R., Di Tacchio, M., et al. (2022). Phenotypic determinism and stochasticity in antibody repertoires of clonally expanded plasma cells. Proc. Natl. Acad. Sci. U. S. A. 119, e2113766119.

104. Correa, A., Trajtenberg, F., Obal, G., Pritsch, O., Dighiero, G., Oppezzo, P., and Buschiazzo, A. (2013). Structure of a human IgA1 Fab fragment at 1.55 Å resolution: potential effect of the constant domains on antigen-affinity modulation. Acta Crystallogr. D Biol. Crystallogr. 69, 388–397.

105. Pan, S., Manabe, N., and Yamaguchi, Y. (2021). 3D Structures of IgA, IgM, and Components. Int. J. Mol. Sci. 22. 10.3390/ijms222312776.

106. Otten, M.A., and van Egmond, M. (2004). The Fc receptor for IgA (FcαRI, CD89). Immunology Letters 92, 23–31. 10.1016/j.imlet.2003.11.018.

107. Wegman, A.D., Fang, H., Rothman, A.L., Thomas, S.J., Endy, T.P., McCracken, M.K., Currier, J.R., Friberg, H., Gromowski, G.D., and Waickman, A.T. (2021). Monomeric IgA Antagonizes IgG-Mediated Enhancement of DENV Infection. Front. Immunol. 12, 777672.

108. Halstead, S.B. (2014). Dengue Antibody-Dependent Enhancement: Knowns and Unknowns. Microbiol Spectr 2. 10.1128/microbiolspec.AID-0022-2014.

109. Rouers, A., Appanna, R., Chevrier, M., Lum, J., Lau, M.C., Tan, L., Loy, T., Tay, A., Sethi, R., Sathiakumar, D., et al. (2021). CD27CD38 plasmablasts are activated B cells of mixed origin with distinct function. iScience 24, 102482.

110. Monteiro, R.C., and Van De Winkel, J.G.J. (2003). IgA Fc receptors. Annu. Rev. Immunol. 21, 177–204.

111. Chevailler, A., Monteiro, R.C., Kubagawa, H., and Cooper, M.D. (1989). Immunofluorescence analysis of IgA binding by human mononuclear cells in blood and lymphoid tissue. J. Immunol. 142, 2244–2249.

112. Maliszewski, C.R., Shen, L., and Fanger, M.W. (1985). The expression of receptors for IgA on human monocytes and calcitriol-treated HL-60 cells. J. Immunol. 135, 3878–3881.

113. Monteiro, R.C., Kubagawa, H., and Cooper, M.D. (1990). Cellular distribution, regulation, and biochemical nature of an Fc alpha receptor in humans. J. Exp. Med. 171, 597–613.

114. Durbin, A.P., Vargas, M.J., Wanionek, K., Hammond, S.N., Gordon, A., Rocha, C., Balmaseda, A., and Harris, E. (2008). Phenotyping of peripheral blood mononuclear cells during acute dengue illness demonstrates infection and increased activation of monocytes in severe cases compared to classic dengue fever. Virology 376, 429–435.

115. Jessie, K., Fong, M.Y., Devi, S., Lam, S.K., and Wong, K.T. (2004). Localization of dengue virus in naturally infected human tissues, by immunohistochemistry and in situ hybridization. J. Infect. Dis. 189, 1411–1418.

116. Kyle, J.L., Beatty, P.R., and Harris, E. (2007). Dengue virus infects macrophages and dendritic cells in a mouse model of infection. J. Infect. Dis. 195, 1808–1817.

117. Kou, Z., Quinn, M., Chen, H., Rodrigo, W.W.S.I., Rose, R.C., Schlesinger, J.J., and Jin, X. (2008). Monocytes, but not T or B cells, are the principal target cells for dengue virus (DV) infection among human peripheral blood mononuclear cells. J. Med. Virol. 80, 134–146.

118. Balsitis, S.J., Coloma, J., Castro, G., Alava, A., Flores, D., McKerrow, J.H., Beatty, P.R., and Harris, E. (2009). Tropism of dengue virus in mice and humans defined by viral nonstructural protein 3-specific immunostaining. Am. J. Trop. Med. Hyg. 80, 416–424.

119. Pasquier, B., Launay, P., Kanamaru, Y., Moura, I.C., Pfirsch, S., Ruffié, C., Hénin, D., Benhamou, M., Pretolani, M., Blank, U., et al. (2005). Identification of FcalphaRI as an inhibitory receptor that controls inflammation: dual role of FcRgamma ITAM. Immunity 22, 31–42.

120. Breedveld, A., and van Egmond, M. (2019). IgA and FcαRI: Pathological Roles and Therapeutic Opportunities. Front. Immunol. 10, 553.

121. Dengue vaccine: WHO position paper, September 2018 - Recommendations (2019). Vaccine 37, 4848–4849.

122. Cassetti, M.C., Durbin, A., Harris, E., Rico-Hesse, R., Roehrig, J., Rothman, A., Whitehead, S., Natarajan, R., and Laughlin, C. (2010). Report of an NIAID workshop on dengue animal models. Vaccine 28, 4229–4234.

123. Chen, R.E., and Diamond, M.S. (2020). Dengue mouse models for evaluating pathogenesis and countermeasures. Curr. Opin. Virol. 43, 50–58.

124. Clark, K.B., Onlamoon, N., Hsiao, H.-M., Perng, G.C., and Villinger, F. (2013). Can non-human primates serve as models for investigating dengue disease pathogenesis? Front. Microbiol. 4, 305.

125. van Egmond, M., van Vuuren, A.J., Morton, H.C., van Spriel, A.B., Shen, L., Hofhuis, F.M., Saito, T., Mayadas, T.N., Verbeek, J.S., and van de Winkel, J.G. (1999). Human immunoglobulin A receptor (FcalphaRI, CD89) function in transgenic mice requires both FcR gamma chain and CR3 (CD11b/CD18). Blood 93, 4387–4394.

126. Davis, C.W., Nguyen, H.-Y., Hanna, S.L., Sánchez, M.D., Doms, R.W., and Pierson, T.C. (2006). West Nile virus discriminates between DC-SIGN and DC-SIGNR for cellular attachment and infection. J. Virol. 80, 1290–1301.

127. Guthmiller, J.J., Dugan, H.L., Neu, K.E., Lan, L.Y.-L., and Wilson, P.C. (2019). An Efficient Method to Generate Monoclonal Antibodies from Human B Cells. Methods Mol. Biol. 1904, 109–145.

128. Pierson, T.C., Sánchez, M.D., Puffer, B.A., Ahmed, A.A., Geiss, B.J., Valentine, L.E., Altamura, L.A., Diamond, M.S., and Doms, R.W. (2006). A rapid and quantitative assay for measuring antibody-mediated neutralization of West Nile virus infection. Virology 346, 53–65.

129. Ansarah-Sobrinho, C., Nelson, S., Jost, C.A., Whitehead, S.S., and Pierson, T.C. (2008). Temperature-dependent production of pseudoinfectious dengue reporter virus particles by complementation. Virology 381, 67–74.

130. Dowd, K.A., Mukherjee, S., Kuhn, R.J., and Pierson, T.C. (2014). Combined effects of the structural heterogeneity and dynamics of flaviviruses on antibody recognition. J. Virol. 88, 11726– 11737.

131. Dowd, K.A., DeMaso, C.R., Pelc, R.S., Speer, S.D., Smith, A.R.Y., Goo, L., Platt, D.J., Mascola, J.R., Graham, B.S., Mulligan, M.J., et al. (2016). Broadly Neutralizing Activity of Zika Virus-Immune Sera Identifies a Single Viral Serotype. Cell Rep. 16, 1485–1491.

132. Mattia, K., Puffer, B.A., Williams, K.L., Gonzalez, R., Murray, M., Sluzas, E., Pagano, D., Ajith, S., Bower, M., Berdougo, E., et al. (2011). Dengue reporter virus particles for measuring neutralizing antibodies against each of the four dengue serotypes. PLoS One 6, e27252.

133. Walker, L.M., Huber, M., Doores, K.J., Falkowska, E., Pejchal, R., Julien, J.-P., Wang, S.-K., Ramos, A., Chan-Hui, P.-Y., Moyle, M., et al. (2011). Broad neutralization coverage of HIV by multiple highly potent antibodies. Nature 477, 466–470.

134. Oliphant, T., Nybakken, G.E., Engle, M., Xu, Q., Nelson, C.A., Sukupolvi-Petty, S., Marri, A., Lachmi, B.-E., Olshevsky, U., Fremont, D.H., et al. (2006). Antibody recognition and neutralization determinants on domains I and II of West Nile Virus envelope protein. J. Virol. 80, 12149–12159.

135. Zhao, H., Fernandez, E., Dowd, K.A., Speer, S.D., Platt, D.J., Gorman, M.J., Govero, J., Nelson, C.A., Pierson, T.C., Diamond, M.S., et al. (2016). Structural Basis of Zika Virus-Specific Antibody Protection. Cell 166, 1016–1027. 10.1016/j.cell.2016.07.020.

136. Vogt, M.R., Moesker, B., Goudsmit, J., Jongeneelen, M., Kyle Austin, S., Oliphant, T., Nelson, S., Pierson, T.C., Wilschut, J., Throsby, M., et al. (2009). Human Monoclonal Antibodies against West Nile Virus Induced by Natural Infection Neutralize at a Postattachment Step. Journal of Virology 83, 6494–6507. 10.1128/jvi.00286-09.

